# Mitochondrial perturbation in the intestine causes microbiota-dependent injury and gene signatures discriminative of inflammatory disease

**DOI:** 10.1101/2023.07.27.549844

**Authors:** Elisabeth Urbauer, Doriane Aguanno, Nora Mindermann, Hélène Omer, Amira Metwaly, Tina Krammel, Tim Faro, Marianne Remke, Sandra Reitmeier, Stefanie Bärthel, Johannes Kersting, Zihua Huang, Feng Xian, Manuela Schmidt, Dieter Saur, Samuel Huber, Bärbel Stecher, Markus List, David Gómez-Varela, Katja Steiger, Matthieu Allez, Eva Rath, Dirk Haller

## Abstract

Mitochondrial dysfunction is associated with inflammatory bowel diseases (IBD). To understand how microbial-metabolic circuits contribute to intestinal tissue injury, we disrupt mitochondrial function in the epithelium by deleting heat shock protein 60 (Hsp60^Δ/ΔIEC^). While metabolic perturbation causes self-resolving tissue injury, regeneration is disrupted in the absence of aryl hydrocarbon receptor (Hsp60^Δ/ΔIEC^;AhR^-/-^) or IL-10 (Hsp60^Δ/ΔIEC^;Il10^-/-^) leading to IBD-like pathology. Injury is absent in the distal colon of germ-free (GF) Hsp60^Δ/ΔIEC^ mice, highlighting bacterial control of metabolic injury. Selective colonization of GF Hsp60^Δ/ΔIEC^ mice with the synthetic community OMM^12^ confirms consistent expansion of metabolically-flexible *Bacteroides* spp. across all models and mono-colonization with *B. caecimuris* recapitulates injury. Transcriptional profiling of metabolically-impaired epithelium identifies gene signatures, including *Ido1*, *Nos2*, and *Duox2*, distinguishing active from inactive tissue inflammation in 343 resected samples from Crohn’s disease patients. In conclusion, mitochondrial perturbation of the epithelium causes microbiota-dependent tissue injury and discriminative inflammatory gene profiles relevant for IBD.

Graphical Abstract:
Control of metabolic injury by microbial signals.

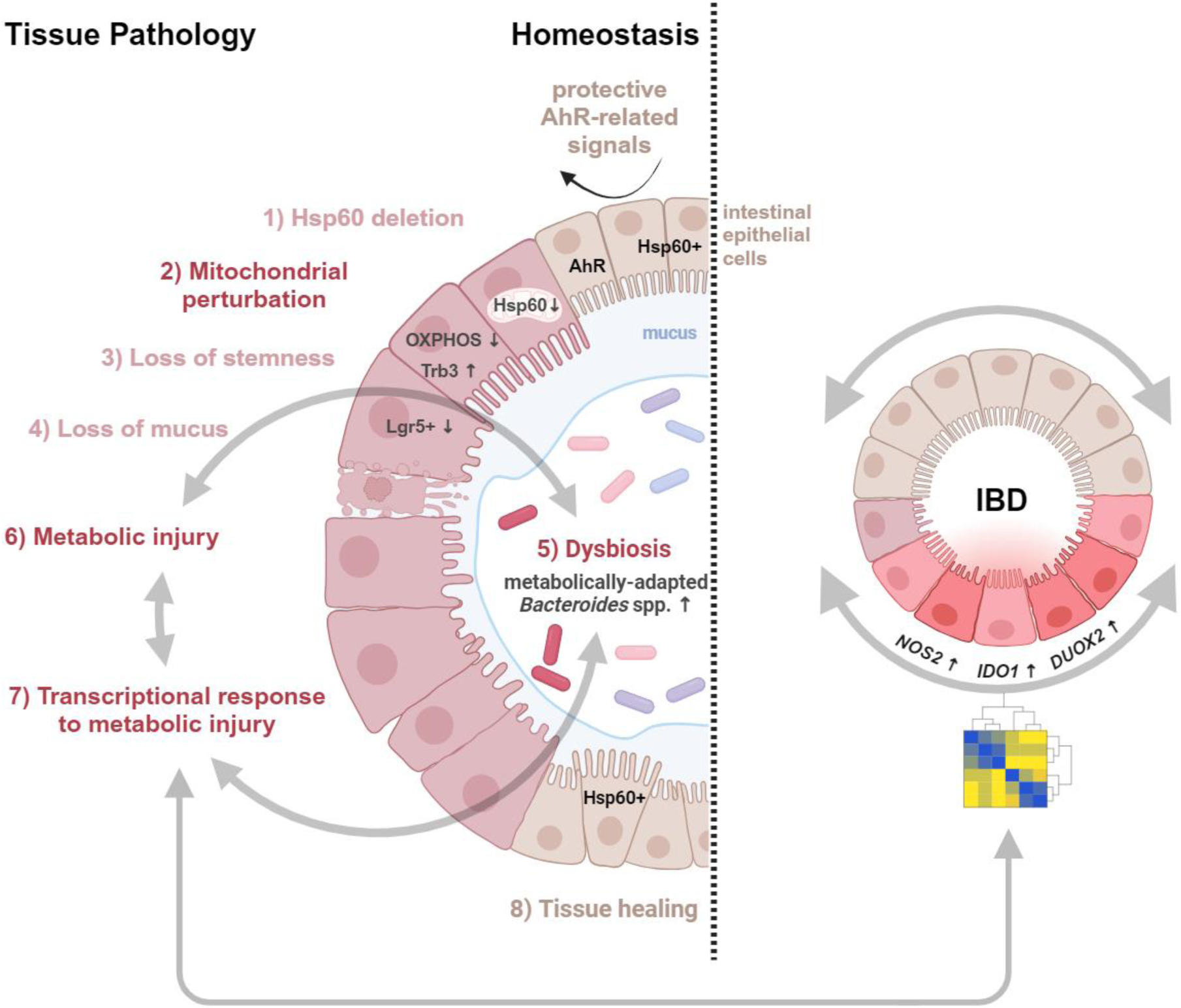

1. Mitochondrial perturbation of the intestinal epithelium induces tissue injury
2. Loss of IL-10 and AhR-related host mechanisms accelerate injury and inflammation
3. Mitochondrial dysfunction induces dysbiosis and expansion of *Bacteroides* spp.
4. Metabolic injury gene signature discriminates inflamed *vs.* non-inflamed IBD samples

## Introduction

The intestinal epithelium is exposed to a diverse and complex microbial milieu at the mucosal interface. A breach in epithelial barrier integrity is associated with inflammatory pathologies^1^ and necessitates a fine-tuned balance of epithelial renewal for healing processes. In particular, the high energy demand of the inflamed mucosa requires mitochondria as central coordinators of metabolic circuits between the host and the microbiome, adding a new aspect to the “energy deficiency” hypothesis^2^. Mitochondria control intestinal epithelial cell (IEC) metabolism, including oxidative phosphorylation (OXPHOS), and influence IEC fate and properties^3^. A cornerstone in this homeostatic regulation is the mitochondrial unfolded protein response (MT-UPR), a conserved signaling pathway triggered upon protein aggregation or misfolding in the mitochondrial matrix^4^. A key molecule of this autoregulatory mechanism is the mitochondrial heat shock protein 60 (Hsp60), a chaperone controlling the refolding of imported as well as misfolded proteins^3^. While activated MT-UPR signaling, and subsequently increased Hsp60 expression, aims to restore homeostasis during acute stress by controlling epithelial self-renewal^3^, its sustained activation is associated with disease pathogenesis^5,6^.

Microbiome-host mutualism is required for intestinal homeostasis, and its disruption contributes to the initiation and progression of diseases^7^, including inflammatory bowel diseases (IBD) ^8^, graft-versus-host disease^9^, and type 2 diabetes^10^. Among the > 240 identified IBD genetic risk loci, genes associated with mitochondrial function point towards its implication in IBD^11,12^. In line, we previously showed that conditional deletion of Hsp60 in the murine small intestine induces mitochondrial impairment, leading to a loss of stemness and epithelial injury^13^. Accordingly, we demonstrated that activated MT-UPR signaling is linked to IBD-related inflammation in patients^14^. While loss of the mitochondrial protein prohibitin 1 triggers mitochondrial impairment in Paneth cells and subsequent ileitis in mice^15^, mitochondrial dysfunction in the intestinal stem cell (ISC) niche predicts disease recurrence in Crohn’s disease patients^16^. Even though the involvement of the microbiome in IBD has been extensively studied, data on the link between mitochondrial dysfunction and microbial composition is scarce. An intriguing new concept suggests that metabolic reprogramming of IECs contributes to microbial dysbiosis^17^, highlighting the bi-directional cross-talk of the microbiome and the host. We hypothesize that mitochondria are involved in sensing dietary and microbial metabolites, allowing the epithelium to functionally adapt to the luminal intestinal environment. While cellular receptors, such as Toll-like receptors^18^ and aryl hydrocarbon receptor (AhR) ^19^, are involved in controlling IEC homeostasis and mitochondrial metabolism, the molecular mechanisms of mitochondrial signal integration remain elusive. We postulate that failure of mitochondrial function at the intestinal interface and disruption of microbial-metabolic circuits contribute to dysbiosis and chronic intestinal inflammation.

To investigate microbe-host cross-talks and their contribution to metabolic injury as a novel hypothesis for disturbed epithelial regeneration, we induce mitochondrial dysfunction by deleting Hsp60 specifically in murine IECs. We show that mitochondrial impairment causes metabolic injury, a cell-autonomous mechanism of tissue wounding, which accelerates microbiota-dependent IBD-like colonic pathologies in the absence of AhR or Interleukin (IL)-10. Remarkably, mitochondrial perturbation leads to the dysbiotic adaption of metabolically-flexible *Bacteroides caecimuris* causing temporally- and spatially-controlled tissue injury. Finally, we identify a murine gene signature of metabolic injury, which discriminates tissue inflammation in a cohort of Crohn’s disease (CD) patients. Conclusively, these data indicate that mitochondria are a cornerstone in regulating cellular metabolism and function and that loss of microbial-metabolic circuits disrupts intestinal tissue homeostasis and accelerates IBD-like pathologies.

## Results

### Mitochondrial impairment accelerates IL-10 related IBD pathology

Mitochondrial function regulates stem cell homeostasis, and disruption of metabolic circuits in the intestinal epithelium is an emerging mechanism in the development of recurrent inflammation^3^. To address the functional consequences of mitochondrial dysfunction in IECs, we deleted *Hsp60* (Hsp60^Δ/ΔIEC^) in mice housed under specific pathogen-free (SPF) conditions, causing transitory weight loss (Figure 1A, B). Gene expression analysis confirmed transient *Hsp60* deletion (Figure S1A) and activation of MT-UPR signaling, measured *via* the marker gene *tribbles pseudokinase 3* (*Trb3*) (Figure S1B) in the colon.

**Figure 1:**
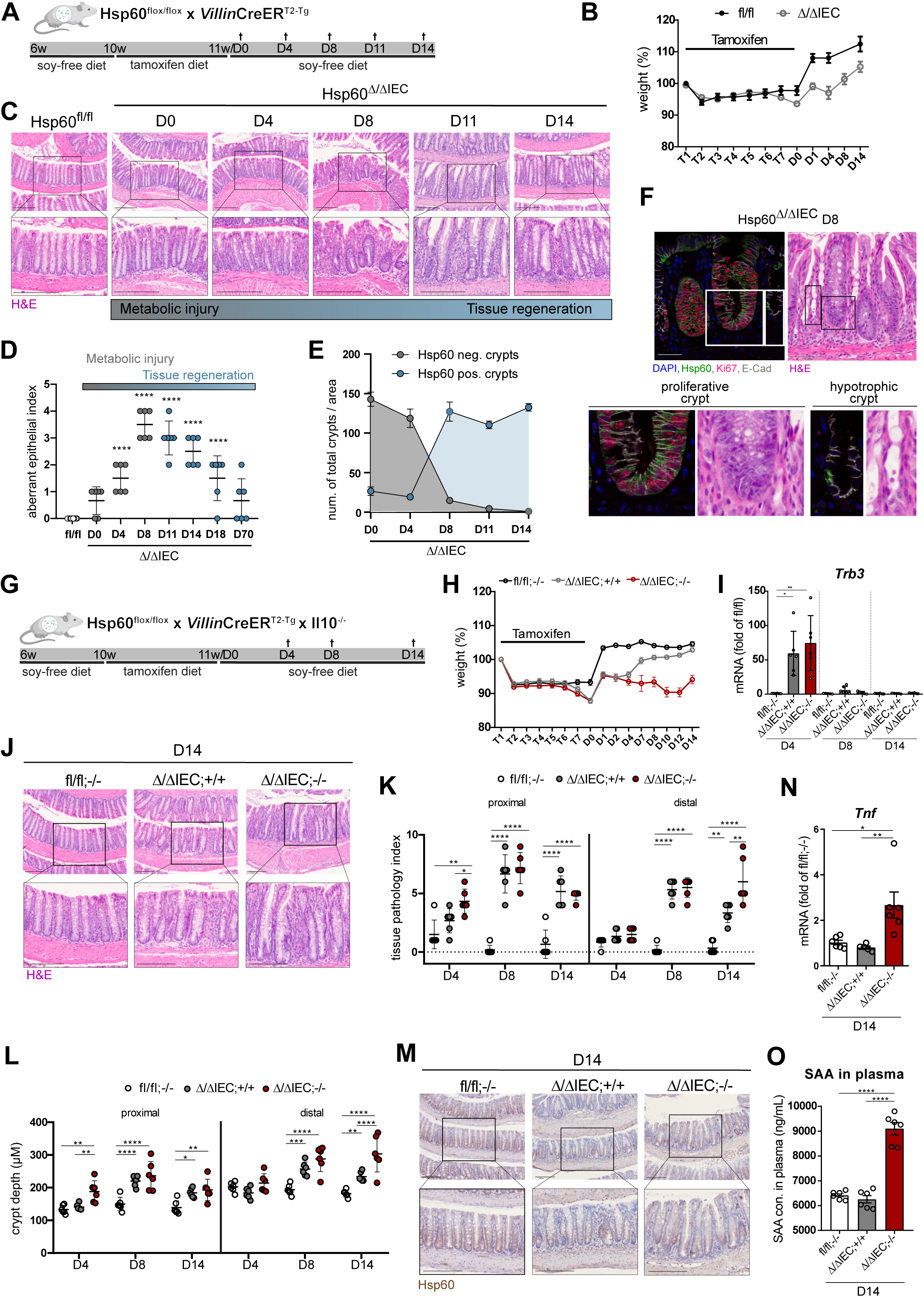
Metabolic injury accelerates IL-10 related IBD pathology. **A,** Experimental setup. Hsp60^fl/fl^ x *Villin*CreER^T2-Tg^ mice (Hsp60^Δ/ΔIEC^; Ctrl: Hsp60^fl/fl^) were fed a soy-free diet for 4 weeks, followed by a tamoxifen diet at the age of 10 weeks for 7 days to induce Hsp60 deletion in IECs. Mice were sampled 0, 4, 8, 11, and 14 days after tamoxifen feeding and full Hsp60 deletion. **B,** Relative weight curve of Hsp60^Δ/ΔIEC^ and Hsp60^fl/fl^ mice from the start of tamoxifen feeding until sampling time point. **C,** Representative images of H&E staining of colonic tissue of Hsp60^Δ/ΔIEC^ and Hsp60^fl/fl^ mice from day 0 to 14. Scale bars: 200µm. **D,** H&E staining of colonic tissue of Hsp60^Δ/ΔIEC^ and Hsp60^fl/fl^ mice from day 0 to 70 were histologically scored for an aberrant epithelium. **E,** Quantification of Hsp60-negative and -positive crypts in Hsp60^Δ/ΔIEC^ mice from day 0 to 14. **F,** Representative immunofluorescent co-staining of Hsp60 (green), Ki67 (red), E-Cadherin (grey), and DAPI (blue) as well as H&E staining of different crypt morphologies at day 8 in colonic tissue of Hsp60^Δ/ΔIEC^ mice. Scale bars: 50µm. **G,** Experimental setup. Hsp60^fl/fl^ x *Villin*CreER^T2-Tg^ x Il10^-/-^ mice (Hsp60^Δ/ΔIEC^;Il10^-/-^; Ctrl: Hsp60^Δ/ΔIEC^;Il10^+/+^, Hsp60^fl/fl^;Il10^-/-^) were fed a soy-free diet for 4 weeks, followed by a tamoxifen diet at the age of 10 weeks for 7 days to induce Hsp60 deletion. Mice were sampled 4, 8, and 14 days after tamoxifen feeding and full Hsp60 deletion. **H,** Relative weight curve of Hsp60^Δ/ΔIEC^;Il10^-/-^, Hsp60^Δ/ΔIEC^;Il10^+/+^ and Hsp60^fl/fl^;Il10^-/-^ mice from the start of tamoxifen feeding until sampling time point. **I,** qPCR of the MT-UPR marker gene *Trb3* using mRNA of whole colonic tissue at day 4, 8, and 14 in Hsp60^Δ/ΔIEC^;Il10^-/-^ (Ctrl: Hsp60^Δ/ΔIEC^;Il10^+/+^, Hsp60^fl/fl^;Il10^-/-^) mice. **J,** Representative images of H&E staining of colonic tissue of Hsp60^Δ/ΔIEC^;Il10^-/-^ (Ctrl: Hsp60^Δ/ΔIEC^;Il10^+/+^, Hsp60^fl/fl^;Il10^-/-^) mice at day 14. Scale bars: 200µm. **K, L,** H&E stainings of colonic tissue (proximal *vs*. distal) of Hsp60^Δ/ΔIEC^;Il10^-/-^ (Ctrl: Hsp60^Δ/ΔIEC^;Il10^+/+^, Hsp60^fl/fl^;Il10^-/-^) mice from day 4, 8, and 14 were (**K**) histologically scored for inflammation and (**L**) crypt length was measured. **M,** Representative images of immunohistochemistry staining of Hsp60 using colonic tissue of Hsp60^Δ/ΔIEC^;Il10^-/-^ (Ctrl: Hsp60^Δ/ΔIEC^;Il10^+/+^, Hsp60^fl/fl^;Il10^-/-^) mice at day 14. The brown color shows specific Hsp60 signal, and nuclei are stained with hematoxylin. Scale bars: 200µm. **N, O,** (**N**) qPCR of *Tnf* using mRNA of whole colonic tissue from day 14 and (**O**) ELISA of SAA using serum from day 14 of Hsp60^Δ/ΔIEC^;Il10^-/-^ (Ctrl: Hsp60^Δ/ΔIEC^;Il10^+/+^, Hsp60^fl/fl^;Il10^-/-^) mice. Data are represented by (**B, H**) mean ± sem or (**D, E, I, K, L, N, O**) mean ± s.d. of six biological replicates. (**D, E, I, K, L, O**) P-values were calculated by ANOVA followed by pairwise comparison testing (Tukey). (**N**) P-values were calculated by Kruskal-Wallis test followed by pairwise comparison testing (Dunn). *P<0.05, **P<0.01, ***P<0.001, ****P<0.0001. D: day; Hsp60: heat shock protein 60; IECs: intestinal epithelial cells; H&E: hematoxylin and eosin; DAPI: 4′,6-diamidino-2-phenylindole; E-Cad: E-Cadherin; MT-UPR: mitochondrial unfolded protein response; Trb3: tribbles pseudokinase 3; pos.: positive; neg.: negative; IL10: interleukin-10; Tnf: tumor necrosis factor; SAA: serum amyloid A. See also Figures S1 and S2.

Interestingly, colonic histopathological evaluation revealed a time-dependent restructuration of epithelial architecture following Hsp60 loss (maximal at day 0), characterized by the appearance of aberrant crypts at day 8 (Figure 1C, D; Figure S1C). This metabolic injury phenotype was followed by regenerative responses and a regain of Hsp60-positive crypts (Figure 1E; Figure S1D), leading to tissue restitution and healing at day 14 in Hsp60^Δ/ΔIEC^ animals (Figure 1C, D; Figure S1C). *Ex vivo* Hsp60-deletion in intestinal organoids also led to the activation of MT-UPR signaling (*Trb3*), reduced stemness (*Lgr5*) and reduced ATP production (Figure S1E-I), confirming the effects of Hsp60 loss on epithelial homeostasis at the cellular level.

At the molecular level, microarray analysis of Hsp60-deficient colonocytes confirmed MT-UPR induction, including the regulatory *activating transcription factor 5* (*Atf5*), and highlighted that mitochondrial dysfunction is associated with metabolic reprogramming (reduction in OXPHOS, increase in glycolysis) (Figure S1J). Remarkably, while inducible overexpression of *Atf5* in the intestinal epithelium (Atf5^IEC^; Figure S1K) confirmed induction of MT-UPR signaling (*Trb3)*, Atf5^IEC^ transgenic mice did not develop colonic tissue injury (Figure S1L-Q), suggesting that functional impairment of mitochondria, rather than their stress response, is responsible for tissue injury.

To better understand the profound morphological changes in the colon caused by metabolic disruption, we characterized key features of the epithelial barrier. Regarding the regenerative capacity, we observed highly proliferative crypt nodules (Ki67-positive and bromodeoxyuridine (BrdU)-positive crypts; Figure S2A-D), which were constituted of Hsp60-positive cells (Figure 1F). The latter presumably originated from stem cells escaping Hsp60 deletion^13^ and contributed to repopulate the epithelium. In parallel, few hypotrophic quiescent crypts, both negative for Hsp60 and the proliferation marker Ki67 (Figure 1F) were observed. Noteworthy, barrier integrity was maintained during metabolic injury, as illustrated by the correct localization of tight junction proteins Zonula occludens-1 (ZO-1) and occludin at the surface epithelium and in crypt nodules (Figure S2E). *Tjp1* and *Ocln* expression levels were even elevated during healing at day 14 (Figure S2F, G). Furthermore, this tissue reorganization was not accompanied by epithelial cell apoptosis (Figure S2H), despite the moderate increase in the inflammation markers *Saa3* and *Tnf* levels (Figure S2I, J).

To better delineate the contribution of mitochondrial impairment of the epithelium and metabolic injury on the onset of IBD-related inflammation and pathology, we crossed Hsp60^Δ/ΔIEC^ mice with IL-10-deficient mice (Hsp60^Δ/ΔIEC^;Il10^-/-^; Figure 1G). In contrast to Hsp60^Δ/ΔIEC^;Il10^+/+^ mice, Hsp60^Δ/ΔIEC^;Il10^-/-^ mice did not recover from weight loss (Figure 1H). Of note, IL-10 deficiency did not further impact genetically-driven activation of *Trb3* (Figure 1I). Most remarkably, at day 14, Hsp60^Δ/ΔIEC^;Il10^-/-^ mice displayed a disrupted epithelial reconstitution in the distal colon (Figure 1J), which was corroborated by an elevated tissue pathology index (Figure 1K) and elongated crypts (Figure 1L), demonstrating a disrupted regenerative response to metabolic injury in the absence of IL-10. Of note, the regain of Hsp60 was independent of tissue regeneration (Figure 1M). In line with tissue injury and an impaired barrier integrity, illustrated by disrupted tight junctional complexes (Figure S2E), gene expression of *Tnf* and serum protein levels of serum amyloid A (SAA) were increased (Figure 1N, O). Since Hsp60^fl/fl^;Il10^-/-^ control mice did not develop colitis at such an early time point, these data provide further evidence that metabolic injury accelerates the onset or recurrence of chronic inflammation followed by a disturbed epithelial regeneration in an IBD susceptible host.

### Metabolic injury drives dysbiotic changes in the intestinal microbiota

Since microbial dysbiosis is a cornerstone in IBD pathogenesis, we analyzed the intestinal microbiome in the course of metabolic injury. 16S rRNA amplicon sequencing of colonic content revealed that metabolic injury was associated with rapid but transient changes in microbiota composition, including a drop in species richness (Figure 2A), as well as changes in the community profile (β-diversity) (Figure 2B). Analysis of differentially abundant taxa revealed six significantly altered genera, including *Bacteroides* spp. (Figure 2C). Interestingly, *Bacteroides* spp. showed the highest overall relative abundance and were transiently increased during metabolic injury, illustrating the rapid adaption of the microbiota to disturbances in IEC metabolism. Shallow shotgun metagenomics precisely revealed the presence of *B. caecimuris, B. uniformis,* and *Phocaeicola vulgatus* (Figure 2D). Impaired mucus barrier is a hallmark of IBD pathogenesis and, concordantly, our results showed that the number of mucus-filled goblet cells (GCs) was strongly reduced in Hsp60^Δ/ΔIEC^ mice from day 4 onwards (Figure S3A, B), suggesting that disrupted epithelial-microbial interactions start prior to visible tissue injury.

**Figure 2:**
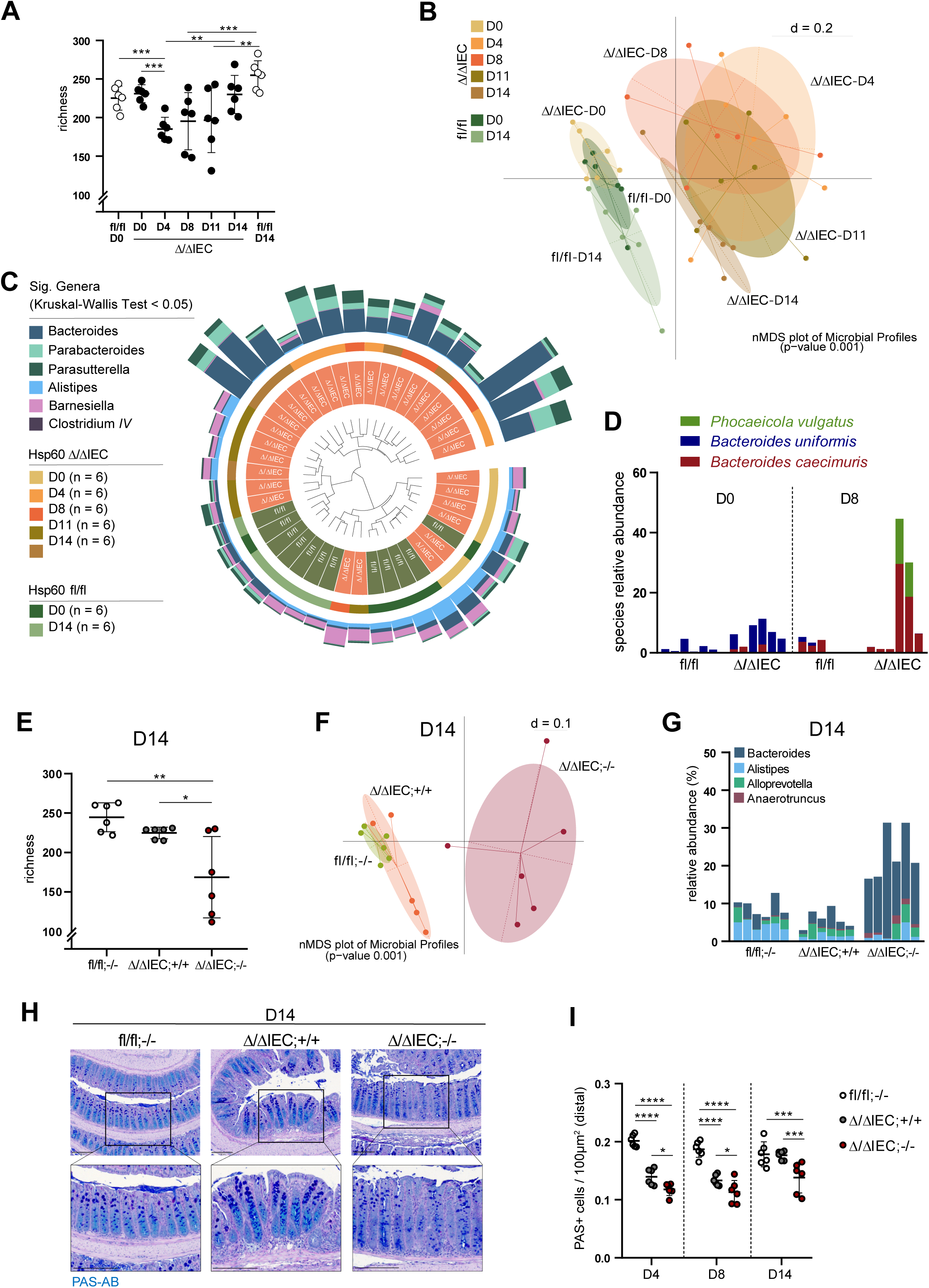
Metabolic injury drives dysbiotic changes in the intestinal microbiota. **A,** Species richness in colonic content of Hsp60^Δ/ΔIEC^ and Hsp60^fl/fl^ mice from day 0 to 14. **B,** β-diversity depicted as nMDS plot of microbial profiles of colonic content samples stratified by genotype (Hsp60^Δ/ΔIEC^ and Hsp60^fl/fl^ mice) and time point (day 0, 4, 8, 11, 14). **C,** Relative abundance of significantly enriched genera of the colonic microbiota in Hsp60^Δ/ΔIEC^ and Hsp60^fl/fl^ mice from day 0 to 14. The dendrogram depicts similarities between microbiota profiles based on generalized UniFrac distances between all 42 mice. Individual taxonomic composition of significantly altered genera is shown as stacked bar plots around the dendrogram. **D,** Shallow shotgun metagenomics of colonic microbiota in Hsp60^Δ/ΔIEC^ and Hsp60^fl/fl^ mice at day 0 and 8. Species relative abundance of the three detected *Bacteroides* species, *B. caecimuris*, *B. uniformis,* and *Phocaeicola vulgatus,* is shown. **E,** Species richness in colonic content of Hsp60^Δ/ΔIEC^;Il10^-/-^ (Ctrl: Hsp60^Δ/ΔIEC^;Il10^+/+^, Hsp60^fl/fl^;Il10^-/-^) mice at day 14. **F,** β-diversity depicted as nMDS plot of microbial profiles of colonic content stratified by genotype (Hsp60^Δ/ΔIEC^;Il10^-/-^, Hsp60^Δ/ΔIEC^;Il10^+/+^, Hsp60^fl/fl^;Il10^-/-^) at day 14. **G,** Relative abundance of significantly enriched genera in Hsp60^Δ/ΔIEC^;Il10^-/-^ (Ctrl: Hsp60^Δ/ΔIEC^;Il10^+/+^, Hsp60^fl/fl^;Il10^-/-^) mice at day 14. **H,** Representative images of PAS-AB staining in colonic tissue of Hsp60^Δ/ΔIEC^;Il10^-/-^ (Ctrl: Hsp60^Δ/ΔIEC^;Il10^+/+^ and Hsp60^fl/fl^;Il10^-/-^) mice at day 14. Acidic mucins are stained in cyan, neutral mucins in dark blue. Nuclei are stained with hematoxylin. Scale bars: 200µm. **I,** Quantification of PAS-positive cells per 100µm^2^ in distal colonic tissue of Hsp60^Δ/ΔIEC^;Il10^-/-^ (Ctrl: Hsp60^Δ/ΔIEC^;Il10^+/+^, Hsp60^fl/fl^;Il10^-/-^) mice at day 4, 8, and 14. (**A, E, I**) Data are represented by mean ± s.d. of six biological replicates. P-values were calculated by ANOVA followed by pairwise comparison testing (Tukey). *P<0.05, **P<0.01, ***P<0.001, ****P<0.0001. (**B, G**) Kruskal-Wallis test was used to select significant genera (P<0.05). D: day; Hsp60: heat shock protein 60; IECs: intestinal epithelial cells; IL10: interleukin-10; PAS-AB: Alcian blue/Periodicacid-Schiff. See also Figure S3.

While these microbiota changes were transient in Hsp60^Δ/ΔIEC^ mice, alterations in species richness and β-diversity were more pronounced at day 8 (Figure S3C-E) and continued at day 14 in Hsp60^Δ/ΔIEC^;Il10^-/-^ mice (Figure 2E, F). Remarkably, the increased relative abundance of *Bacteroides* spp. persisted in Hsp60^Δ/ΔIEC^;Il10^-/-^ mice (Figure 2G), suggesting that sustained changes in the luminal milieu favor the selective expansion of these bacteria. Consistent with a disrupted regenerative response, Hsp60^Δ/ΔIEC^;Il10^-/-^ mice showed a persistent reduction in GC numbers (Figure 2H, I).

### AhR signaling is a key pathway in regulating metabolic injury

The host sensor AhR is a major regulator in intestinal homeostasis. AhR activation has been demonstrated to be protective against intestinal inflammation in various studies^20,21^ and its deficiency contributes to IBD pathogenesis^21,22^. Moreover, AhR participates in maintaining the ISC niche against damage^23^ and has recently been shown to modulate mitochondrial function^19^. In this context, we aimed to investigate the involvement of AhR in modulating injury and regenerative processes. Therefore we crossed full-body AhR-knockout mice (*AhR^-/-^*) with Hsp60^Δ/ΔIEC^ mice (Hsp60^Δ/ΔIEC^;AhR^-/-^, Figure 3A). SPF-housed Hsp60^Δ/ΔIEC^;AhR^-/-^ mice displayed a severe weight loss (Figure 3B) and reduced survival reaching abortion criteria between day 0 and day 2 (Figure S4A). Histopathological evaluation at day 0 showed a severe loss of epithelial architecture in the ileum of Hsp60^Δ/ΔIEC^;AhR^-/-^ mice (Figure S4B, C), demonstrating a dramatically accelerated tissue pathology leading to death in the absence of AhR signaling.

**Figure 3:**
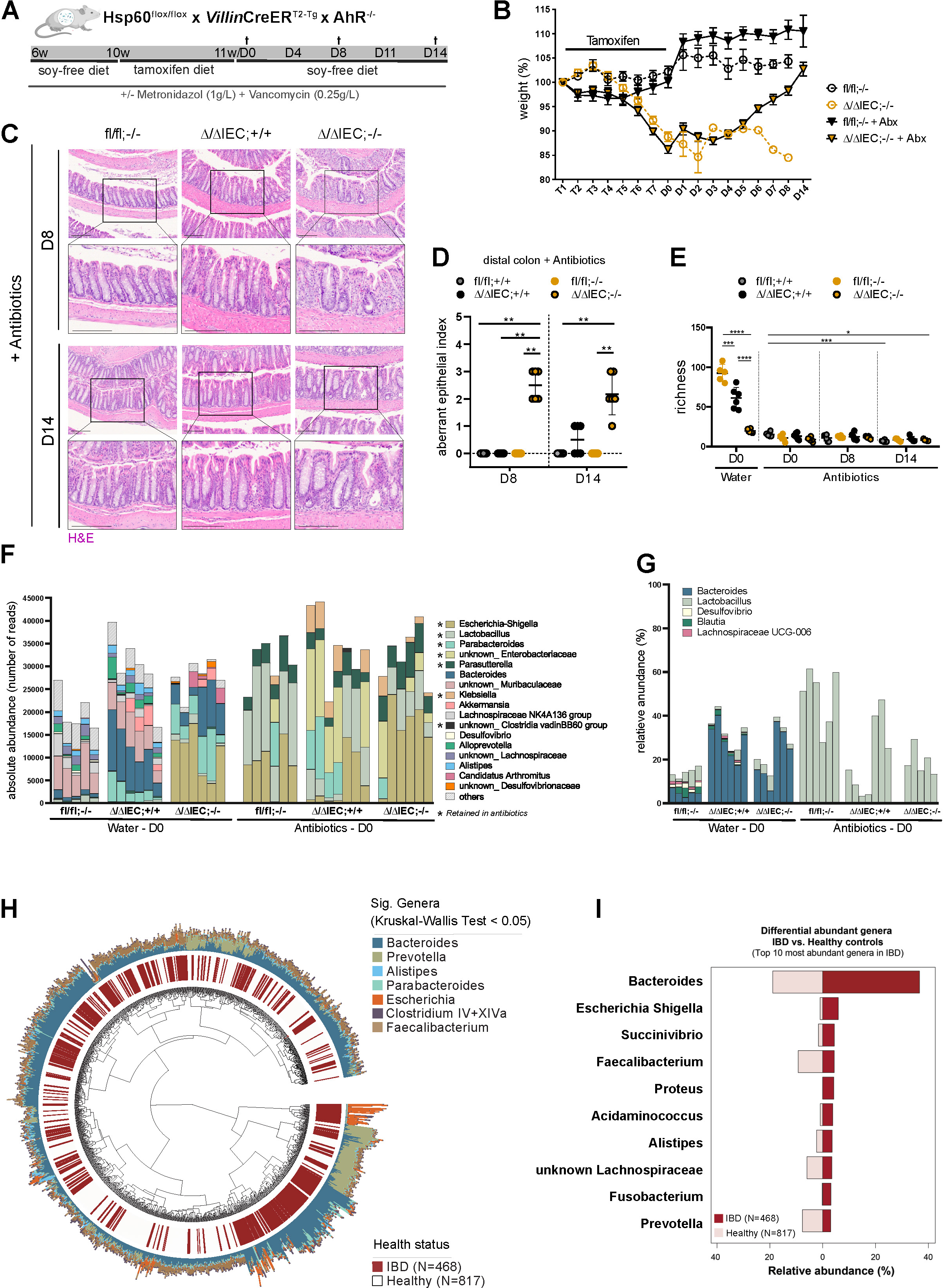
AhR deficiency during metabolic injury is lethal and eradication of *Bacteroides* spp. by antibiotics rescues survival. **A,** Experimental setup. Hsp60^fl/fl^ x *Villin*CreER^T2-Tg^ x AhR^-/-^ mice (Hsp60^Δ/ΔIEC^;AhR^-/-^; Ctrl: Hsp60^Δ/ΔIEC^;AhR^+/+^, Hsp60^fl/fl^;AhR^-/-^) were fed a soy-free diet for 4 weeks, followed by a tamoxifen diet at the age of 10 weeks for 7 days to induce Hsp60 deletion. In addition, mice were treated with antibiotics (Abx; Metronidazole (1g/L) and Vancomycin (0.25g/L)) from 6 weeks on. Mice were sampled at day 0, 8, and 14 after tamoxifen feeding and full Hsp60 deletion. In addition to a control cohort (water) sampled at day 0, animals exposed to water were either sampled at day 8 (Ctrl: Hsp60^Δ/ΔIEC^;AhR^+/+^, Hsp60^fl/fl^;AhR^-/-^) or when reaching abortion criteria for Hsp60^Δ/ΔIEC^;AhR^-/-^ mice. **B,** Relative weight curve of Hsp60^Δ/ΔIEC^;AhR^-/-^ mice treated with or without Abx (Ctrl: Hsp60^fl/fl^;AhR^-/-^) from the start of tamoxifen feeding until sampling time point. **C,** Representative images of H&E staining of distal colonic tissue of Hsp60^Δ/ΔIEC^;AhR^-/-^ mice (Ctrl: Hsp60^Δ/ΔIEC^;AhR^+/+^, Hsp60^fl/fl^;AhR^-/-^) at day 8 and 14 treated with Abx. Scale bars: 200µm. **D,** H&E stainings of distal colonic tissue of Hsp60^Δ/ΔIEC^;AhR^-/-^ mice (Ctrl: Hsp60^Δ/ΔIEC^;AhR^+/+^, Hsp60^fl/fl^;AhR^-/-^) at day 8 and 14 treated with Abx were histologically scored for an aberrant epithelium. **E,** Species richness in colonic content of Hsp60^Δ/ΔIEC^;AhR^-/-^ mice (Ctrl: Hsp60^Δ/ΔIEC^;AhR^+/+^, Hsp60^fl/fl^;AhR^-/-^) treated with (day 0, 8 and 14) or without antibiotics (day 0). **F,** Absolute abundance of different genera of the colonic microbiota in Hsp60^Δ/ΔIEC^;AhR^-/-^ mice (Ctrl: Hsp60^Δ/ΔIEC^;AhR^+/+^, Hsp60^fl/fl^;AhR^-/-^) treated with or without Abx (day 0). Genera still present upon Abx treatment are indicated by an asterisk. **G**, Relative abundance of selected genera of the colonic microbiota in Hsp60^Δ/ΔIEC^;AhR^-/-^ mice (Ctrl: Hsp60^Δ/ΔIEC^;AhR^+/+^, Hsp60^fl/fl^;AhR^-/-^) treated with or without Abx (day 0). Only genera significantly differing in relative abundances (Kruskal-Wallis, Dunn, P<0.05) between water-exposed Hsp60^fl/fl^;AhR^-/-^ and Hsp60^Δ/ΔIEC^;AhR^+/+^ groups on day 0 are displayed. **H,** Relative abundance of significantly enriched genera of the fecal microbiota in IBD patients (N=468) and healthy controls (N=817). The dendrogram depicts similarities between microbiota profiles based on generalized UniFrac distances between all 1285 individuals. Individual taxonomic composition at the genus level is shown as stacked bar plots around the dendrogram in addition to health status (red: IBD patients). **I,** Relative abundance of the top ten differential abundant genera in IBD patients (N=468) versus healthy controls (N=817). Data are represented by (**B**) mean ± sem or (**D, E**) mean ± s.d. of six biological replicates. (**E**) P-values were calculated by ANOVA followed by pairwise comparison testing (Tukey). (**D**) P-values were calculated by Kruskal-Wallis test followed by pairwise comparison testing (Dunn). *P<0.05, **P<0.01, ***P<0.001, ****P<0.0001. (**G, H**) Kruskal-Wallis test was used to select significant genera (P<0.05). *P<0.05, **P<0.01, ***P<0.001, ****P<0.0001. D: day; Hsp60: heat shock protein 60; IECs: intestinal epithelial cells; AhR: aryl hydrocarbon receptor; Abx: antibiotics; H&E: hematoxylin and eosin; IBD: inflammatory bowel diseases. See also Figure S4.

To further elucidate the protective role of AhR, we addressed the involvement of IL-22, a major downstream signaling pathway of AhR. Since IL-22 was increased in colonic tissue during metabolic injury (Figure S4D), we crossed full-body IL-22 receptor subunit-α1-knockout mice (*IL22ra1*^-/-^) with Hsp60^Δ/ΔIEC^ mice (Hsp60^Δ/ΔIEC^;Il22ra1^-/-^) (Figure S4E). Interestingly, the absence of the IL-22 receptor did not affect weight development during metabolic injury (Figure S4F). In line, histological evaluation demonstrated no difference in epithelial aberration with or without IL22ra1 (Figure S4G-I), indicating that this cytokine pathway is not involved in modulating metabolic injury and regeneration.

Interestingly, while exposure to antibiotics abrogated the lethal phenotype of Hsp60^Δ/ΔIEC^;AhR^-/-^ mice at early stages (Figure 3A, B; Figure S4A), tissue injury was retained at day 8 and day 14, unlike in control littermates (Figure 3C, D). Species richness was strongly reduced in Hsp60^Δ/ΔIEC^;AhR^-/-^ mice (most likely due to diarrhea), compared to water controls, and, as expected, the administration of antibiotics maintained low community richness in all groups (Figure 3E). Strikingly, the sustained increase in absolute (Figure 3F) and relative abundance (Figure 3G) of *Bacteroides* spp., observed in Hsp60^Δ/ΔIEC^;AhR^-/-^ and Hsp60^Δ/ΔIEC^;AhR^+/+^ mice (water groups), was totally abrogated in antibiotic-treated animals (Figure 3F, G). These data clearly support the hypothesis that *Bacteroides* spp. contribute to the development and severity of intestinal injury. In addition, bacterial profiling of fecal samples from 468 IBD patients^24,25^ showed an increased relative abundance of *Bacteroides* spp. associated with intestinal inflammation compared to healthy controls (Figure 3H, I), underlining the importance of these bacteria in IBD.

### Bacterial control of metabolic injury

Given the described dysbiosis occurring during metabolic injury and the striking attenuating effect of antibiotics, we further evaluated the causative role of bacteria in metabolic injury by generating germ-free (GF) Hsp60^Δ/ΔIEC^ mice (Figure 4A). Compared to SPF housing, weight reduction in GF Hsp60^Δ/ΔIEC^ mice was mild (Figure 4B), despite activated MT-UPR signaling (*Trb3*) (Figure 4C) and a similar Hsp60 regain in the crypt epithelium in the absence of bacteria (Figure 4D, E; Figure S5A) compared to previous results in the presence of bacteria. This is in line with our finding that induction of MT-UPR signaling in Atf5^IEC^ mice alone did not translate into tissue injury. Histopathological evaluation and analysis of *Tnf* and *Saa3* gene expression confirmed the absence of metabolic injury and inflammation in the middle and distal colon (Figure 4F-H; Figure S5B, C), supporting the hypothesis that bacteria are involved in regulating metabolic injury. Both proximal colon and ileum largely retained an aberrant epithelial architecture (Figure 4G, H; Figure S5D-F), suggesting that bacteria-independent mechanisms drive the injury phenotype in these tissue segments. Despite the absence of tissue pathology under GF conditions, Ki67 and BrdU staining of the epithelium showed increased colonic proliferation in Hsp60^Δ/ΔIEC^ mice compared to control animals (Figure S5G-J), however to a lesser extent compared to SPF-housed mice (Figure S2A-D). The analysis of the mucus barrier in Hsp60^Δ/ΔIEC^ mice showed only a mild reduction in GC numbers compared to control littermates (Figure S5K-L), unlike previously seen under SPF housing. Junctional complexes of the epithelium were not changed at protein or transcriptional levels (Figure S5M-O), indicating a preserved intestinal barrier during metabolic injury in the absence of bacteria.

**Figure 4:**
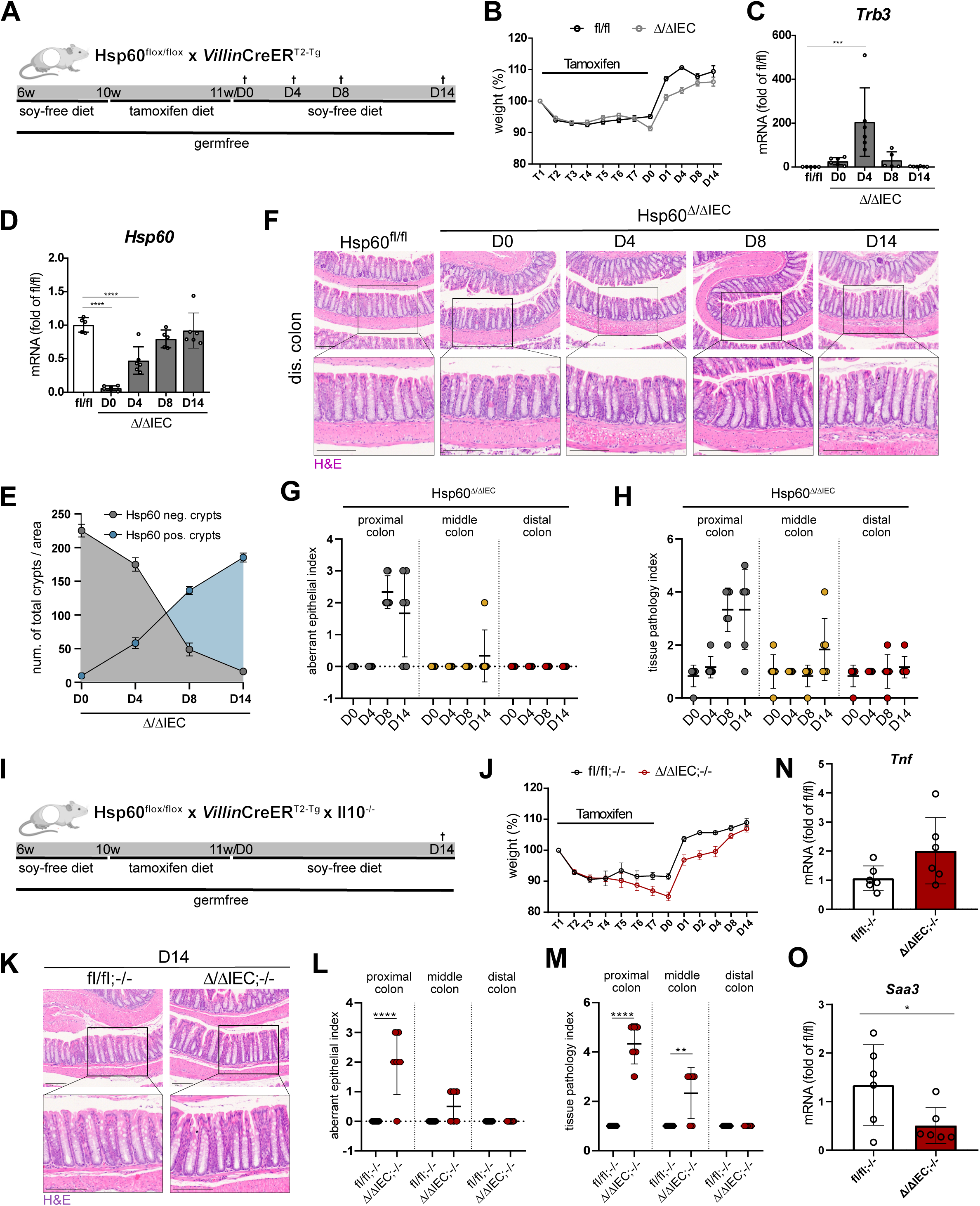
Bacteria control metabolic injury. **A,** Experimental setup. GF Hsp60^fl/fl^ x *Villin*CreER^T2-Tg^ mice (Hsp60^Δ/ΔIEC^; Ctrl: Hsp60^fl/fl^) were fed a soy-free diet for 4 weeks, followed by a tamoxifen diet at the age of 10 weeks for 7 days to induce Hsp60 deletion in IECs. Mice were sampled 0, 4, 8, and 14 days after tamoxifen feeding and full Hsp60 deletion. **B,** Relative weight curve of GF Hsp60^Δ/ΔIEC^ and Hsp60^fl/fl^ mice from the start of tamoxifen feeding until sampling time point. **C, D,** qPCR of (**C**) the MT-UPR marker gene *Trb3* and (**D**) the mitochondrial chaperone *Hsp60* using mRNA of whole colonic tissue from day 0 to 14 in GF Hsp60^Δ/ΔIEC^ and Hsp60^fl/fl^ mice. **E,** Quantification of Hsp60-negative and -positive crypts in GF Hsp60^Δ/ΔIEC^ mice from day 0 to 14. **F,** Representative images of H&E staining of distal colonic tissue of GF Hsp60^Δ/ΔIEC^ and Hsp60^fl/fl^ mice from day 0 to 14. Scale bars: 200µm. **G, H,** H&E stainings of colonic tissue of GF Hsp60^Δ/ΔIEC^ mice from day 0 to 14 were histologically scored for (**G**) an aberrant epithelium and (**H**) inflammation. The colonic swiss role was divided into three parts (proximal, middle and distal) and scored separately. **I,** Experimental setup. GF Hsp60^fl/fl^ x *Villin*CreER^T2-Tg^ x Il10^-/-^ mice (Hsp60^Δ/ΔIEC^;Il10^-/-^; Ctrl: Hsp60^fl/fl^;Il10^-/-^) were fed a soy-free diet for 4 weeks, followed by a tamoxifen diet at the age of 10 weeks for 7 days to induce Hsp60 deletion. Mice were sampled 14 days after tamoxifen feeding and full Hsp60 deletion. **J,** Relative weight curve of GF Hsp60^Δ/ΔIEC^;Il10^-/-^ (Ctrl: Hsp60^fl/fl^;Il10^-/-^) mice from the start of tamoxifen feeding until sampling time point. **K,** Representative images of H&E staining of distal colonic tissue of GF Hsp60^Δ/ΔIEC^;Il10^-/-^ (Ctrl: Hsp60^fl/fl^;Il10^-/-^) mice at day 14. Scale bars: 200µm. **L, M,** H&E stainings of colonic tissue of GF Hsp60^Δ/ΔIEC^;Il10^-/-^ (Ctrl: Hsp60^fl/fl^;Il10^-/-^) mice from day 14 were histologically scored for (**L**) an aberrant epithelium and (**M**) inflammation. The colonic swiss role was divided into three parts (proximal, middle and distal) and scored separately. **N,O,** qPCR of the inflammatory marker genes (**N**) *Tnf* and (**O**) *Saa3* using mRNA of whole colonic tissue from day 0 to 14 in GF Hsp60^Δ/ΔIEC^;Il10^-/-^ and Hsp60^fl/fl^;Il10^-/-^ mice. Data are represented by (**B, J**) mean ± sem or (**C, D, G, H, L-O**) mean ± s.d. of five to six biological replicates. P-values were calculated by (**C, D**) ANOVA followed by pairwise comparison testing (Tukey), (**G, H**) Kruskal-Wallis test followed by pairwise comparison testing (Dunn) or (**L-O**) unpaired t-test. GF: germfree; D: day; Hsp60: heat shock protein 60; IECs: intestinal epithelial cells; H&E: hematoxylin and eosin; MT-UPR: mitochondrial unfolded protein response; pos.: positive; neg.: negative; IL10: interleukin-10; Tnf: Tumor necrosis factor; Trb3: tribbles pseudokinase 3; Saa3: serum amyloid A 3. See also Figure S5.

Similar to Hsp60-deficient mice, and despite the increased inflammatory disease susceptibility, GF Hsp60^Δ/ΔIEC^;Il10^-/-^ mice did not reveal severe weight loss (Figure 4I, J). Importantly, histopathological evaluation confirmed the absence of tissue injury (Figure 4K-M) and inflammation (Figure 4N, O) in the distal colon of GF Hsp60^Δ/ΔIEC^;Il10^-/-^ mice, supporting the pivotal role of bacteria in regulating metabolic injury and chronic inflammatory phenotypes.

### Metabolically-flexible *B. caecimuris* induces colonic injury

Considering the importance of bacteria in controlling injury of the distal colon, we next used the synthetic minimal bacterial community Oligo-Mouse-Microbiota 12 (OMM^12^) ^26,27^. The colonization of GF Hsp60^Δ/ΔIEC^ mice with OMM^12^ (Figure 5A) did not impact Hsp60 loss (Figure 5B; Figure S6A) and MT-UPR activation (*Trb3*) (Figure 5C). Importantly, histopathological analysis showed metabolic injury in the distal colon, including the presence of Hsp60-positive hyperproliferative crypts as well as Hsp60-negative hypotrophic crypts (Figure 5D; Figure S6A). Epithelial aberration, tissue pathology, and crypt depth were elevated (Figure 5E, F; Figure S6B), providing proof-of-concept that simplified bacterial communities are sufficient to induce metabolic injury in the distal colon. 16S rRNA targeted qPCR was used to quantify bacterial abundances of all 12 strains, showing that *B. caecimuris* was elevated from day 4 onwards (Figure 5G), supporting our previous data in complex intestinal bacterial communities of mice and patients.

**Figure 5:**
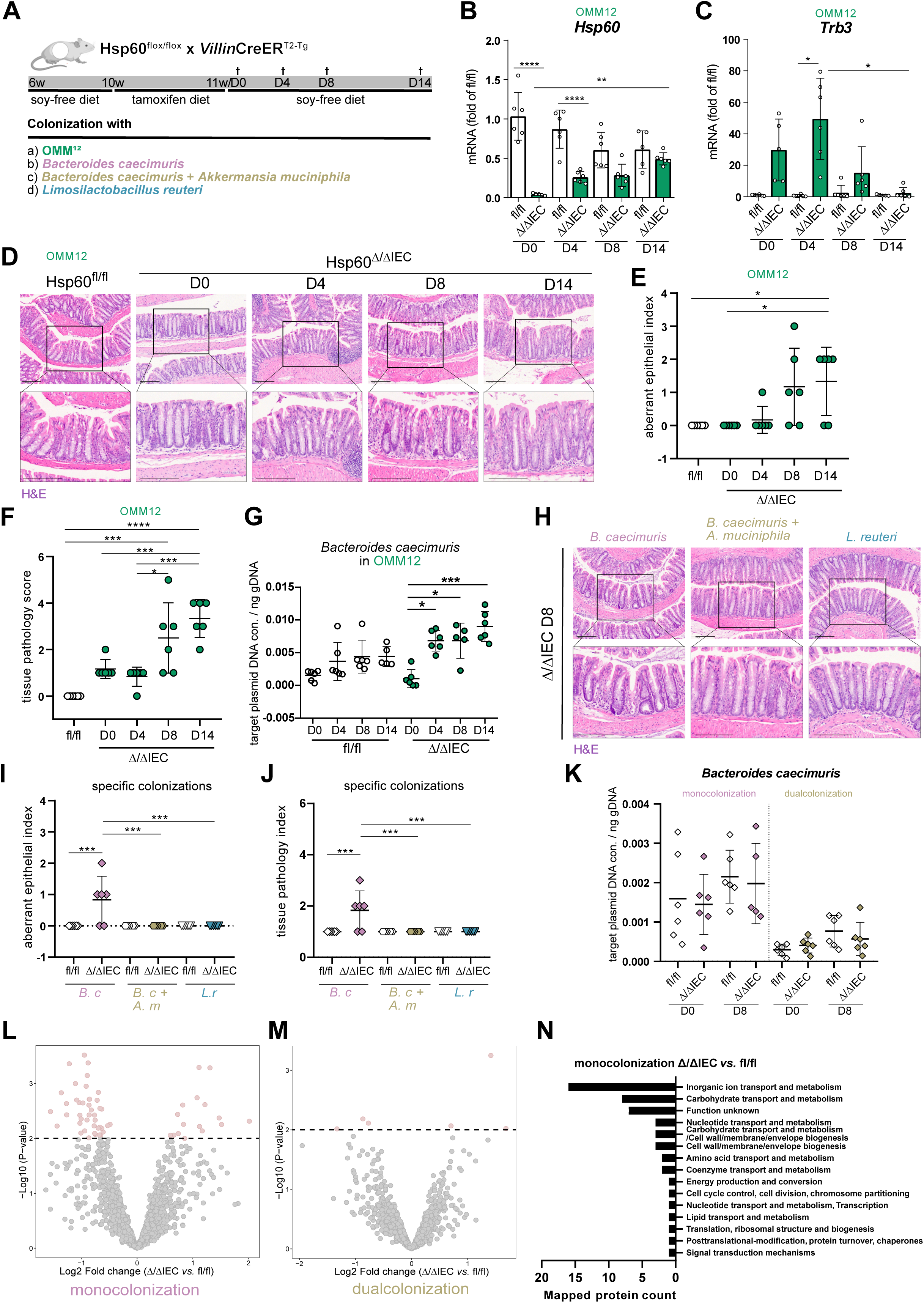
*Bacteroides caecimuris* induces metabolic injury. **A,** Experimental setup. GF Hsp60^fl/fl^ x *Villin*CreER^T2^ mice (Hsp60^Δ/ΔIEC^; Ctrl: Hsp60^fl/fl^) were rectally gavaged twice at the age of 6 weeks with a) the Oligo-Mouse-Microbiota (OMM^12^) synthetic bacterial community, b) *Bacteroides caecimuris*, c) *Bacteroides caecimuris* and *Akkermansia muciniphila* or d) *Limosilactobacillus reuteri*. Mice were fed a soy-free diet for 4 weeks, followed by a tamoxifen diet at the age of 10 weeks for 7 days to induce Hsp60 deletion in IECs. Mice were sampled on different days after tamoxifen feeding and full Hsp60 deletion. **B, C,** qPCR of (**B**) the mitochondrial chaperone *Hsp60* and (**C**) the MT-UPR marker gene *Trb3* using mRNA of whole colonic tissue of GF Hsp60^Δ/ΔIEC^ and Hsp60^fl/fl^ mice colonized with OMM^12^ from day 0 to 14. **D,** Representative images of H&E staining of distal colonic tissue of GF Hsp60^Δ/ΔIEC^ and Hsp60^fl/fl^ mice colonized with OMM^12^ from day 0 to 14. Scale bars: 200µm. **E, F,** H&E stainings of distal colonic tissue of GF Hsp60^Δ/ΔIEC^ and Hsp60^fl/fl^ mice colonized with OMM^12^ were histologically scored for (**E**) an aberrant epithelium and (**F**) inflammation from day 0 to 14. **G,** Abundance of *B. caecimuris* in colonic content of GF Hsp60^Δ/ΔIEC^ and Hsp60^fl/fl^ mice colonized with OMM^12^ quantified by 16S rRNA targeted qPCR from day 0 to 14. **H,** Representative images of H&E staining of distal colonic tissue of GF Hsp60^Δ/ΔIEC^ mice colonized with *B. caecimuris, B. caecimuris* and *A. muciniphila* or *L. reuteri* at day 8. Scale bars: 200µm. **I, J,** H&E stainings of distal colonic tissue of GF Hsp60^Δ/ΔIEC^ mice colonized with *B. caecimuris, B. caecimuris* and *A. muciniphila* or *L. reuteri* were histologically scored for (**I**) an aberrant epithelium and (**J**) inflammation at day 8 after Hsp60 deletion. **K,** Abundance of *B. caecimuris* in colonic content of GF Hsp60^Δ/ΔIEC^ and Hsp60^fl/fl^ mice colonized with *B. caecimuris* or *B. caecimuris* and *A. muciniphila* quantified by 16S rRNA targeted qPCR at day 0 and 8. **L, M,** Volcano plots displaying Log2-fold changes (in Hsp60^Δ/ΔIEC^ over Hsp60^fl/fl^ control mice) of identified protein groups assigned to *B. caecimuris* in, respectively, mono-colonized GF Hsp60^Δ/ΔIEC^ and Hsp60^fl/fl^ mice with (**L**) *B. caecimuris* and (**M**) duo-colonized GF Hsp60^Δ/ΔIEC^ and Hsp60^fl/fl^ mice with *B. caecimuris* and *A. muciniphila*. For both, differentially abundant proteins groups are shown in red (based on P<0.01 Limma R package with moderate t-test). **N,** Counts of protein groups assigned to *B. caecimuris* (and annotated for orthologue function) mapped to COG function categories (see Supplementary Table 1). Data are represented by mean ± s.d. of (**E, F, I, J, K**) six / (**B, C, G, H, I, L, M, N**) five to six biological replicates. P-values were calculated by (**B, I, J, K**) ANOVA followed by pairwise comparison testing (Tukey) or by (**C, E, F, G**) Kruskal-Wallis test followed by pairwise comparison testing (Dunn). *P<0.05, **P<0.01, ***P<0.001, ****P<0.0001. GF: germfree; D: day; Hsp60: heat shock protein 60; IECs: intestinal epithelial cells; MT-UPR: mitochondrial unfolded protein response; OMM^12^: Oligo-Mouse-Microbiota; Trb3: tribbles pseudokinase 3; H&E: hematoxylin and eosin; B.c: *Bacteroides caecimuris*; A.m: *Akkermansia muciniphila*; L.r: *Limosilactobacillus reuteri*; COG: Clusters of Orthologous Genes. See also Figures S6 and S7 and Table S1.

To further dissect the contribution of distinct bacteria in the development of metabolic injury, we mono-colonized GF Hsp60^Δ/ΔIEC^ mice with *B. caecimuris* (Figure 5A) and induced MT-UPR activation through Hsp60 deletion (Figure S6C, D). Strikingly, *B. caecimuris* alone was sufficient to induce metabolic injury in the distal colon, as revealed by the presence of aberrant hyperproliferative crypts (Figure 5H, I; Figure S6E-H) and tissue inflammation (Figure 5J).

Since the abundance of *Akkermansia muciniphila* was slightly elevated during OMM^12^-induced metabolic injury (Figure S7A), we performed a dual-colonization with *B. caecimuris* and *A. muciniphila* (Figure 5A). Interestingly, the presence of *A. muciniphila* reduced the abundance of *B. caecimuris* (Figure 5K) and completely prevented the induction of metabolic injury (Figure 5H-J). Notably, bacterial abundance of *B. caecimuris* and *A. muciniphila* was similar in both genotypes (Figure 5K; Figure S7B).

Mono-colonization of GF Hsp60^Δ/ΔIEC^ mice with *Limosilactobacillus reuteri* (Figure 5A), whose abundance remained unchanged during OMM^12^-induced metabolic injury (Figure S7A), showed again similar *L. reuteri* abundance levels between genotypes (Figure S7C). Importantly, *L. reuteri* did not induce tissue injury in the distal colon (Figure 5H-J).

Finally, we investigated proteome adaptation of *B. caecimuris* during metabolic injury through metaproteomics analysis of colonic content samples of mono- and dual-colonized mice (with, respectively, *B. caecimuris* or *B. caecimuris* and *A. muciniphila*). We identified 60 differentially abundant protein groups (16 being upregulated and 44 being downregulated) in mono-colonized Hsp60^Δ/ΔIEC^ mice compared to Hsp60^fl/fl^ animals (Figure 5L; Figure S6I). Interestingly, and correlating with abrogated injury, the dual-colonization with *A. muciniphila* prevented proteome changes in *B. caecimuris* (Figure 5M). In addition, the annotation of differentially abundant protein groups, mapped to Clusters of Orthologous Groups (COGs) categories, were linked to bacterial metabolism (Figure 5N, Supplementary Table 1), highlighting the ability of *B. caecimuris* to metabolically adapt in response to mitochondrial perturbation and tissue injury.

### Mitochondrial dysfunction in the epithelium defines a transcriptional signature discriminating the inflammation status in IBD

To assess molecular changes during metabolic injury, we performed RNA NanoString^®^ analysis of colonic tissue from SPF and GF Hsp60^Δ/ΔIEC^ mice targeting ∼ 800 genes involved in metabolic processes and immunometabolism. At day 0, 45 genes showed a fold change greater than ±2 under SPF conditions, out of which 10 were mitochondria-related (Figure 6A). Patterns reflecting cellular changes were still present on day 8 (including genes of the solute carrier family and apolipoproteins) but displayed variations, notably with the upregulation of genes involved in oxidative stress as well as immune and antimicrobial responses, including *Duox2*, *Icos*, *Tlr4* and *Nos2*. In contrast, only 10 genes showed a fold change greater than ±2 at day 0 under GF conditions (Figure 6B), indicating that changes in cellular metabolism are dependent on bacterial signals. KEGG pathway analysis of the 45 regulated genes under SPF conditions identified tryptophan metabolism as the most enriched pathway (Figure S8A).

**Figure 6:**
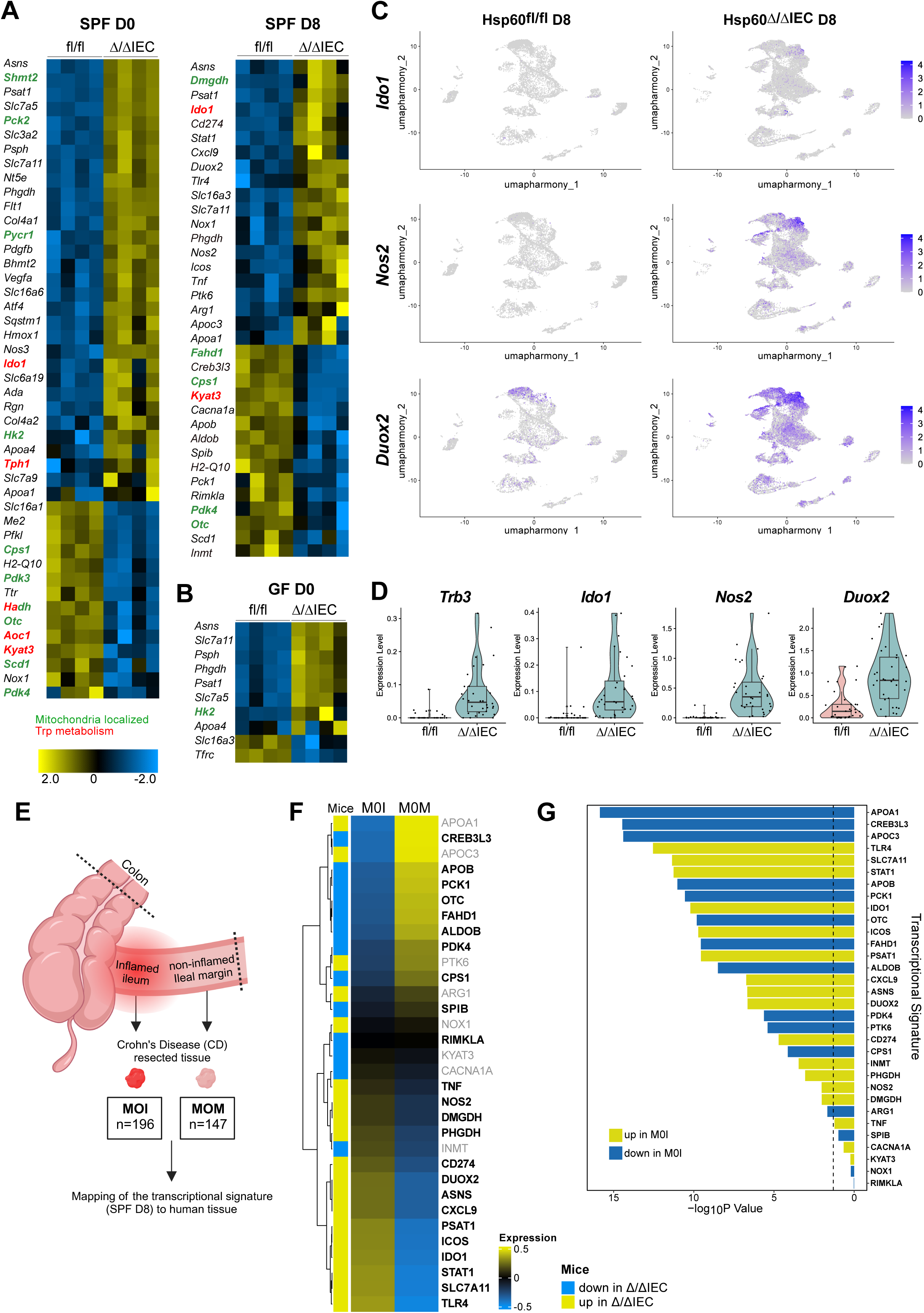
Mitochondrial dysfunction in the epithelium defines a transcriptional signature discriminating the inflammation status in IBD. **A, B,** Heatmap of a NanoString^®^ analysis using a metabolic panel of ∼ 800 genes in mRNA from whole colonic tissue in Hsp60^Δ/ΔIEC^ and Hsp60^fl/fl^ mice under (**A**) SPF or (**B**) GF housing at day 0 and/or 8 after Hsp60 deletion. Depicted are genes with a fold-change greater than ± 2. Genes encoding mitochondria-localized proteins are colored in green, tryptophan metabolism-associated genes in red. **C,** UMAP featuring normalized gene expression levels of, respectively, *Ido1* (top), *Nos2* (middle), *Duox2* (bottom) in Hsp60^Δ/ΔIEC^ and Hsp60^fl/fl^ mice (n=2 per group). **D,** Pseudo-bulk log-normalized transcriptional level of, respectively, *Trb3*, *Ido1*, *Nos2 and Duox2* per condition (Hsp60^Δ/ΔIEC^ and Hsp60^fl/fl^ mice, n=2 per group) across all cell clusters. **E,** Schematic representation of sample origin. Transcriptome data from mucosal biopsies of CD patients (originating from inflamed ileum [M0I] and the non-inflamed Ileal margin [M0M]) were analyzed for transcriptional gene mapping with the metabolic injury signature. **F,** Heatmap of the normalized expression of the 32 human orthologs of the 35 mouse genes identified in SPF-housed animals at D8, based on microarray data of human CD patients. The expression values were gene-wise scaled. Samples from two different locations at the time of surgery were considered: the inflamed ileum [M0I] (n=196) and the ileal margin [M0M] (n=147). The latter served as control samples of uninflamed tissue. Concordant genes between mouse and human are indicated in black and bold while non-concordant genes are indicated in gray. **G,** Boxplot of DEGs between M0M and M0I. 26 of 32 signature genes show significant difference between M0M and M0I groups, and 6 of the signature genes have no significant difference (adjust p value < 0.05). The vertical dashed line represents the cutoff-value of the adjusted p value. (**A, B**) Data include four biological replicates. (**C, D**) Data include two biological replicates. SPF: specific-pathogen-free; GF: germfree; D: day; Hsp60: heat shock protein 60; IECs: intestinal epithelial cells; UMAP: Uniform manifold approximation and projection; CD: Crohn’s Disease; DEGs: differentially expressed genes; Trb3: tribbles pseudokinase 3; Ido1: indoleamine 2, 3-dioxygenase 1; Nos2: nitric oxide synthase 2; Duox2: dual oxidase 2. See also Figures S8 and S9 and Table S2.

Indoleamine 2,3-dioxygenase 1 (Ido1) is the key enzyme involved in the host-related tryptophan-degrading kynurenine pathway, and is upregulated under SPF conditions at day 0 and day 8. Immunofluorescence analysis confirmed the transient increase in Ido1 expression during metabolic injury in the crypts of SPF-housed Hsp60^Δ/ΔIEC^ mice and persistent expression in Hsp60^Δ/ΔIEC^;Il10^-/-^ mice (Figure S8B, upper and lower panel). On the contrary, Ido1 expression was completely absent under GF conditions (Figure S8B, middle panel), highlighting the importance of microbial signals in rewiring host tryptophan metabolism. To better resolve the contribution of Ido1 in tissue injury, we crossed full body Ido1-knockout mice (*Ido1^-/-^*) with Hsp60^Δ/ΔIEC^ mice (Hsp60^Δ/ΔIEC^;Ido1^-/-^) (Figure S8C). Surprisingly, Hsp60^Δ/ΔIEC^;Ido1^-/-^ mice showed no differences in weight loss (Figure S8D), tissue injury (Figure S8E, F) or inflammation (Figure S8G-I) compared to Hsp60^Δ/ΔIEC^;Ido1^+/+^ littermates. Moreover, *Ido1* was induced in intestinal organoids in response to inflammatory cytokines (Tnf and Ifnγ) but not to luminal factors, such as lipopolysaccharides (LPS) or cecal water (Figure S8J, K). Altogether these data clearly demonstrate that Ido1 is not causally involved in the development of metabolic injury, suggesting Ido1 to be regulated as a consequence of host immune responses to metabolic injury.

To better resolve the cell response to metabolic injury, we next applied single-cell RNA-sequencing (scRNA-seq) on samples enriched in IECs from colonic tissue of Hsp60^Δ/ΔIEC^ and Hsp60^fl/fl^ mice at day 8. We identified 16 different cell clusters, including the main IEC subtypes (Figure S9A). The most abundant cell population (cluster 1, Figure S9A) considerably expanded in Hsp60^Δ/ΔIEC^ mice at day 8 (Figure S9B) and it corresponded to highly proliferative cells (Supplementary Table 2). Additional clusters of differentiated cells with specialized functions such as enterocytes (cluster 4 and 6), enteroendocrine cells (EECs, cluster 15) and tuft cells (cluster 12) were reduced in Hsp60^Δ/ΔIEC^ mice (Figure S9B). Smaller clusters, notably cell populations with stromal features supporting epithelial regeneration^28^ (cluster 16) and antimicrobial response (cluster 5, 7 and 11) expanded in metabolically-injured IECs (Figure S9B, Supplementary Table 2). Interestingly, we observed, predominantly in clusters 5, 7 and 11, an upregulation of genes, which were already identified as increased in Hsp60^Δ/ΔIEC^ mice in the Nanostring^®^ analysis, including *Ido1, Nos2* and *Duox2* (Figure 6C). In line, pseudobulk analysis of scRNA-seq populations in Hsp60^Δ/ΔIEC^ mice confirmed increased expression of the MT-UPR marker gene *Trb3* and elevated levels of *Ido1, Nos2* and *Duox2* (Figure 6D) compared to controls, highlighting the occurrence of a murine transcriptional gene signature associated with metabolic injury.

Given the consistency of the transcriptional adaptation in response to metabolic injury, we addressed the human relevance of this metabolic gene signature in CD patients (Figure 6E). Using available transcriptome data from 343 resected tissue specimens of CD patients (204 total) undergoing surgical therapy^29^, we aimed to distinguish between active and inactive disease status based on our metabolic gene signature. We mapped expression patterns of the 32 murine genes (day 8 SPF, Figure 6A) to the corresponding human orthologues in surgically-resected inflamed ileal tissue samples (M0I) and non-inflamed tissue margins (M0M, used as control samples) (Figure 6F). 139 samples were paired (M0I and M0M from the same patient), 57 patients only appeared in M0I and 8 patients only appeared in M0M. 24 out of the 32 genes showed similar expression patterns between mouse and the respective human tissue sections (Figure 6F, highlighted in black and bold). More precisely, 14 genes, which were upregulated in Hsp60^Δ/ΔIEC^ mice (highlighted in yellow in Figure 6F), also showed higher expression levels in M0I *vs*. M0M (Figure 6F). Similarly, 10 genes being downregulated in Hsp60^Δ/ΔIEC^ mice (highlighted in blue in Figure 6F) had concordant lower expression levels in inflamed human tissue (Figure 6F). In contrast, 8 genes had opposite expression patterns between mouse and human tissue (Figure 6F, in gray). The mutual information (non-parametric measure of relevance) of the 24 concordant and 8 discordant genes was 0.13 (p-value of 0.005, based on 10.000 Monte-Carlo permutations), indicating that significantly more genes of the metabolic injury signature were preserved in their relative expression patterns in the CD cohort. Additionally, 26 of 32 signature genes showed significant differences between M0M and M0I (Figure 6G), further emphasizing the relevance of our metabolic injury signature in IBD pathogenesis. Altogether, these data provide evidence that mitochondrial impairment in IECs imprints a transcriptional response signature, which is applicable to IBD and can discriminate between inflamed and non-inflamed mucosa in IBD patients.

## Discussion

Mitochondria are a cornerstone in the control of cellular metabolism and function^6^. We previously demonstrated that conditional deletion of Hsp60 in the small intestinal epithelium initiates stem cell loss and crypt absorption, providing proof-of-concept that mitochondrial perturbation contributes to tissue injury^13^. In the present study, we identify microbial-metabolic circuits as the underlying cause of colonic tissue injury, suggesting a novel mechanism for recurrent inflammation. Noteworthy, we show that metabolically exhausted IECs maintain barrier integrity, despite the regenerative tissue restructuration. In this context, we identify a population of highly proliferative IECs expanding during metabolic injury, which probably repopulate the epithelium. Interestingly, this cluster (*Kcnq1^+^; Kcne3^+^; Cftr^+^*) is enriched in genes encoding chloride and potassium channels shown to be involved in IBD^30,31^. Another small cluster of cells exhibiting characteristics of wound-associated cells also contribute to epithelial restitution^28,32^. On the contrary, in a susceptibility model of IBD-like inflammation, the resolution of metabolic injury is disrupted, leading to disturbed epithelial regeneration and accelerated tissue pathology. This phenotype comprises the loss of GCs and mucus production, as well as sustained microbial dysbiosis, supporting the hypothesis that mitochondrial perturbations contribute to the pathogenesis of chronic inflammation. IBD is characterized by recurrent cycles of wounding and incomplete healing, making the ISC niche and its flexible adjustment of mitochondrial metabolism imperative for tissue regeneration^3^. In line, postoperative recurrence risk of CD patients is linked to mitochondrial dysfunction in the ISC niche^16^. Persistent MT-UPR signaling associated with sustained expression of Hsp60 in the epithelium of IBD-related mouse models and patients^14^ confirms a putative role of mitochondrial perturbation in chronic inflammation. ß-oxidation and subsequent OXPHOS is the primary energy source of bacteria-exposed colonocytes using microbiota-derived SCFA^3,33^. In line with metabolic reprogramming of the Hsp60-deficient epithelium towards glycolysis and reduced ATP production, colonic mitochondrial remodeling and adaptation of energy production are reported to sustain mucosal healing after colitis^34^.

AhR is an environmental sensor playing a critical role in intestinal immunity and barrier function^35^ and studies have described protective effects of AhR activation in colitis mouse models by promoting regulatory profiles in T-cells^36,37^ and IL-22 signaling^38^. Interestingly, IL-22 deficiency does not affect metabolic injury, thus excluding the role of IL-22 as a downstream signal of AhR in response to mitochondrial perturbation and metabolic injury. Loss of AhR in murine IECs is reported to reduce their differentiation capacity leading to impaired barrier function and increased lethality upon infection with *Citrobacter rodentium*^23^. Considering that AhR integrates numerous signals from the host and the enteric bacterial environment, the contribution of this chemoreceptor to intestinal homeostasis and disease susceptibility remains complex. Recent studies clearly demonstrate that AhR contributes to the homeostatic regulation of the ISC niche^23^, supporting the mortal phenotype of Hsp60^Δ/ΔIEC^;AhR^-/-^ mice. Nevertheless, antibiotic treatment of these mice eradicates *Bacteroides* spp. and rescues Hsp60^Δ/ΔIEC^;AhR^-/-^ mice, underlining the essential contribution of distinct bacterial taxa to metabolic injury. Our results in selective transgenic or knock-out mouse models, respectively addressing the contribution of Atf5-driven MT-UPR activation, IL22-signaling or Ido1-related metabolism, demonstrate that isolated pathways do not necessarily control the pathogenesis of tissue injury, emphasizing that metabolic injury arises from complex interplays of metabolic-microbial circuits. Although we cannot completely resolve the contribution of AhR signaling, we postulate that the complex interplay of AhR and bacterial signals orchestrate the regenerative stem cell capacity in a metabolically-exhausted epithelium. Further investigations are required to disentangle the more precise mechanisms of AhR-related signal integration to regulate mitochondrial function.

Metabolic injury in Hsp60^Δ/ΔIEC^ and Hsp60^Δ/ΔIEC^;Il10^-/-^ mice is absent in the distal colon of GF mice, and antibiotic exposure in Hsp60^Δ/ΔIEC^;AhR^-/-^ mice ameliorates the disease severity, highlighting the importance of microbial-metabolic circuits in inflammation-associated tissue injury and their integration by the host. Of note, tissue injury under Hsp60 deficiency is a cell-autonomous process in the small intestine and proximal colon, demonstrating compartmentalized effects of the microbial environment. In line, Hsp60 deficiency in IECs triggers loss of bacterial richness and changes in the microbiota profile, particularly *Bacteroides* spp. expansion. Dysbiosis precedes the onset of tissue injury in Hsp60^Δ/ΔIEC^ mice and persists in Hsp60^Δ/ΔIEC^;Il10^-/-^ mice, suggesting a causal relationship of microbe-host interactions. Selective colonization of GF Hsp60^Δ/ΔIEC^ mice with OMM^12^ recapitulates the injury phenotype and causes absolute *B. caecimuris* expansion. Finally, mono-colonization with *B. caecimuris* induces metabolic injury, demonstrating a causal role of commensal bacteria in initiating epithelial injury in response to mitochondrial perturbation. These data confirm studies reporting gram-negative bacteria, including *Bacteroides*, to be enriched in IBD^39,40^. Interestingly, *A. muciniphila* is a commensal that abrogates metabolic injury in the presence of *B. caecimuris. A. muciniphila* degrades mucins to produce SCFA, in turn regulating mucus thickness and host energy substrates^41^. While healthy controls are associated with beneficial *Akkermansia* spp.^42^, *A. muciniphila* is reportedly reduced in IBD patients^43^. Dual-colonization with *A. muciniphila* inhibits tissue injury and dampens the adaptive metabolic response in *B. caecimuris*, underlining the complex metabolic interplays between OMM^12^ members, as previously published^44^.

Elevated luminal oxygen due to altered colonic epithelial metabolism is a characteristic feature in IBD patients. A recent study shows that enhancing mitochondrial function moderates the disease-associated dysbiosis through reducing oxygen leakage in the lumen^45^. This suggests a hypothetical mechanism for the metabolic injury-associated dysbiosis in Hsp60^Δ/ΔIEC^ and Hsp60^Δ/ΔIEC^;Il10^-/-^ mice. Studies have shown that anaerobes, in particular *Bacteroides* spp., can adapt to O_2_ exposure, similarly to aerobes, to scavenge H_2_O_2_^46^. Our results consistently show a transcriptional induction of genes encoding enzymes involved in oxidative stress and antimicrobial response, such as Duox2 and the inducible nitric oxide synthase (iNOS, encoded by *Nos2*) as well as changes in colonocyte metabolism, notably tryptophan-related pathways. Hence, we postulate that the complex modifications in the epithelial compartment shape luminal environmental conditions favoring the expansion of Bacteroides. Indeed *B. caecimuris* quickly adapts to changes in diet and nutrient sources, as shown in OMM^12^ consortium^47^, and *Bacteroides* spp. evolve into sub-strains ecotypes in the complex murine gut ecosystem^48^, demonstrating their metabolic plasticity. This is in line with our metaproteomic analysis highlighting the metabolic flexibility of *B. caecimuris* in response to mitochondrial perturbation and injury. Further studies would be required to better characterize the metabolic adaptation and potential strain evolution of *Bacteroides* spp. in response to epithelial mitochondrial impairment and their functional consequences on intestinal homeostasis, leading to tissue injury.

Finally, and very importantly for the translational applicability of our findings, we identify a metabolic gene signature in the murine mucosa which is able to discriminate inflamed *versus* non-inflamed tissue sections in CD patients undergoing surgical therapy^29^. This transcriptional pattern is primarily reflecting an adaptive response to the initial mitochondrial impairment in IECs, outlined by the fact that most of the genes are already altered at day 0 after Hsp60 deficiency. The complex regenerative host response at day 8 integrates metabolic adaptation, as well as immune and anti-microbial responses. Our results are in accordance with recently published scRNA-seq data defining a novel IEC population, which expands with active disease in CD patients and is defined by high expression of *LCN2*, *NOS2*, and *DUOX2* (named LND)^49^. Together, these findings strongly support a contribution of mitochondrial dysfunction of the epithelium in IBD pathogenesis.

In mice, the metabolic gene signature relies on the presence of bacteria, supporting a microbial contribution to disease susceptibility in IBD. The use of paired samples from the same resected tissue region allowed us to partially exclude confounding factors, such as drug exposure, disease history, disease location, emphasizing on the presence or absence of inflammation. Hsp60 deficiency in the epithelium causes a similar tissue injury phenotype in ileal and colonic regions of mice, suggesting a universal relevance of the murine signature across different forms of IBD. Nevertheless, one need to consider that the data acquisition in mice and human originated from different tissue regions. It remains to be elucidated to which extent this signature is shared between small and large intestinal disease location, and whether it is representative of IBD or more generally of inflammation.

In summary, we provide evidence that bacterial communities adjust to changes in epithelial metabolism, creating a dysbiotic adaptation of the microbiota in the intestine. What remains elusive and requires further analysis is how metabolic changes in the epithelium select the growth of distinct members of the bacterial community. Here we demonstrate that metabolic flexibility of *B. caecimuris* coincides with metabolic injury, but the explicit contribution of bacterial pathways in these adverse microbial-metabolic circuits remains to be identified. Mitochondrial perturbation and bacterial signals cooperate in the development of metabolic injury, and host-derived metabolic gene signatures discriminate active and inactive inflammatory processes in IBD patients. Thus, targeting specific mitochondrial pathways or microbe-host circuits may yield new therapeutic options in recurrent inflammatory pathologies in the intestine.

## Lead contact statement

Correspondence to Prof. Dr. Dirk Haller, Chair of Nutrition and Immunology, Technical University of Munich, Gregor-Mendel-Str. 2, 85354 Freising, Germany;

## Acknowledgements

Funded by the Deutsche Forschungsgemeinschaft (DFG, German Research Foundation) – project number 395357507 (SFB 1371, Microbiome Signatures) and project number 469152594 (HA 3148/15-1 - Mitochondrial function in the regulation of intestinal tissue regeneration and inflammation-related pathologies). The graphical abstract was created with BioRender.com.

## Authors’ contributions

Conceptualization, E.U., D.A., D.H.; Writing – Original Draft, E.U., D.A., D.H.; Writing – Review & Editing, E.U., D.A., N.M, D.H.; Investigation, E.U., D.A., N.M, T.K., H.O., M.R., F.X.; Methodology, E.U., E.R.; S.B.; Formal Analysis, A.M., S.R., T.F., J.K., Z.H, F.X., M.L., D.G.V.; Funding Acquisition, E.U., E.R., D.H.; Resources, S.H., D.S., B.S., M.L., D.G.V., K.S., M.A.

## STAR Methods

### Key resources table

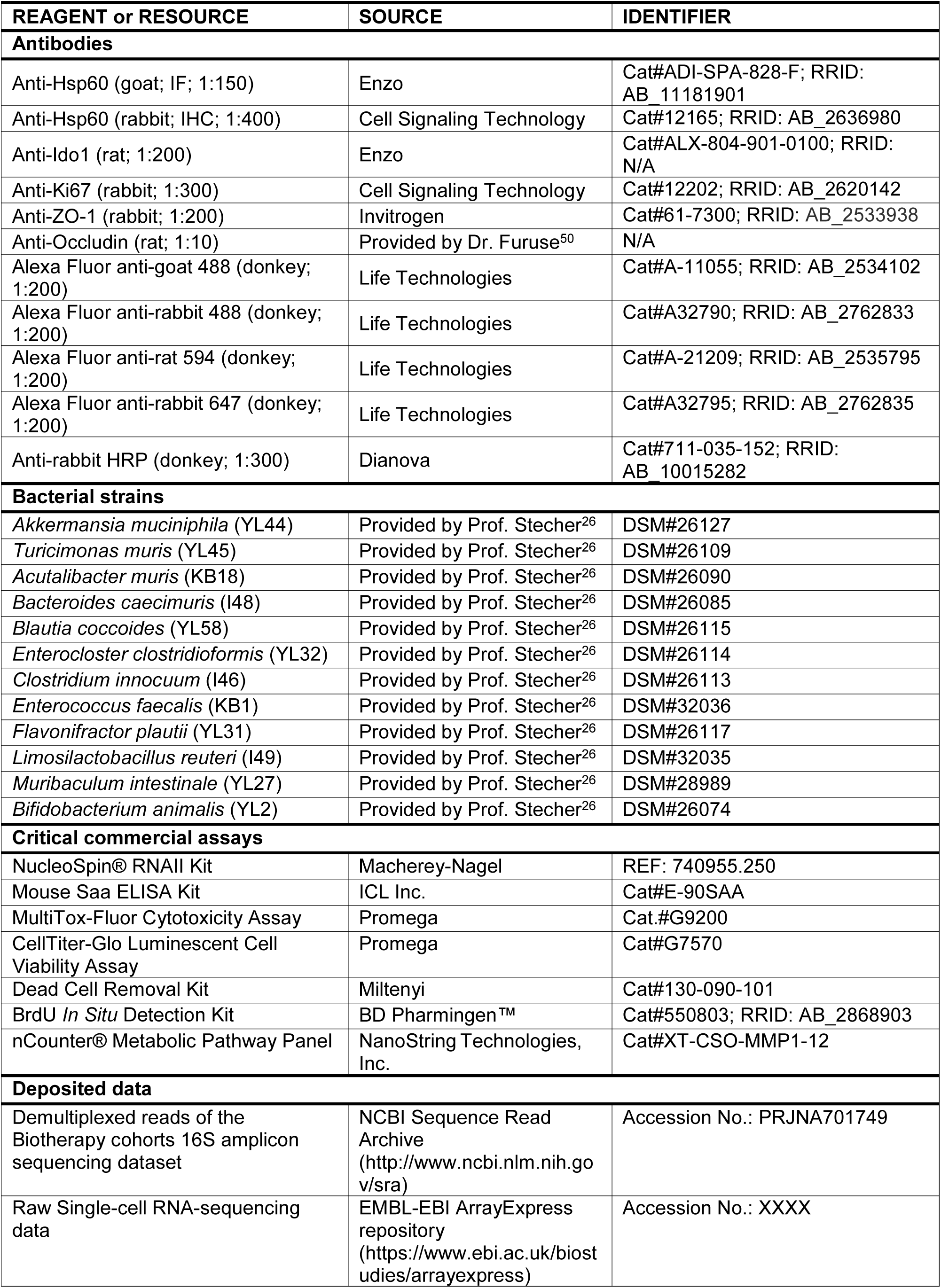
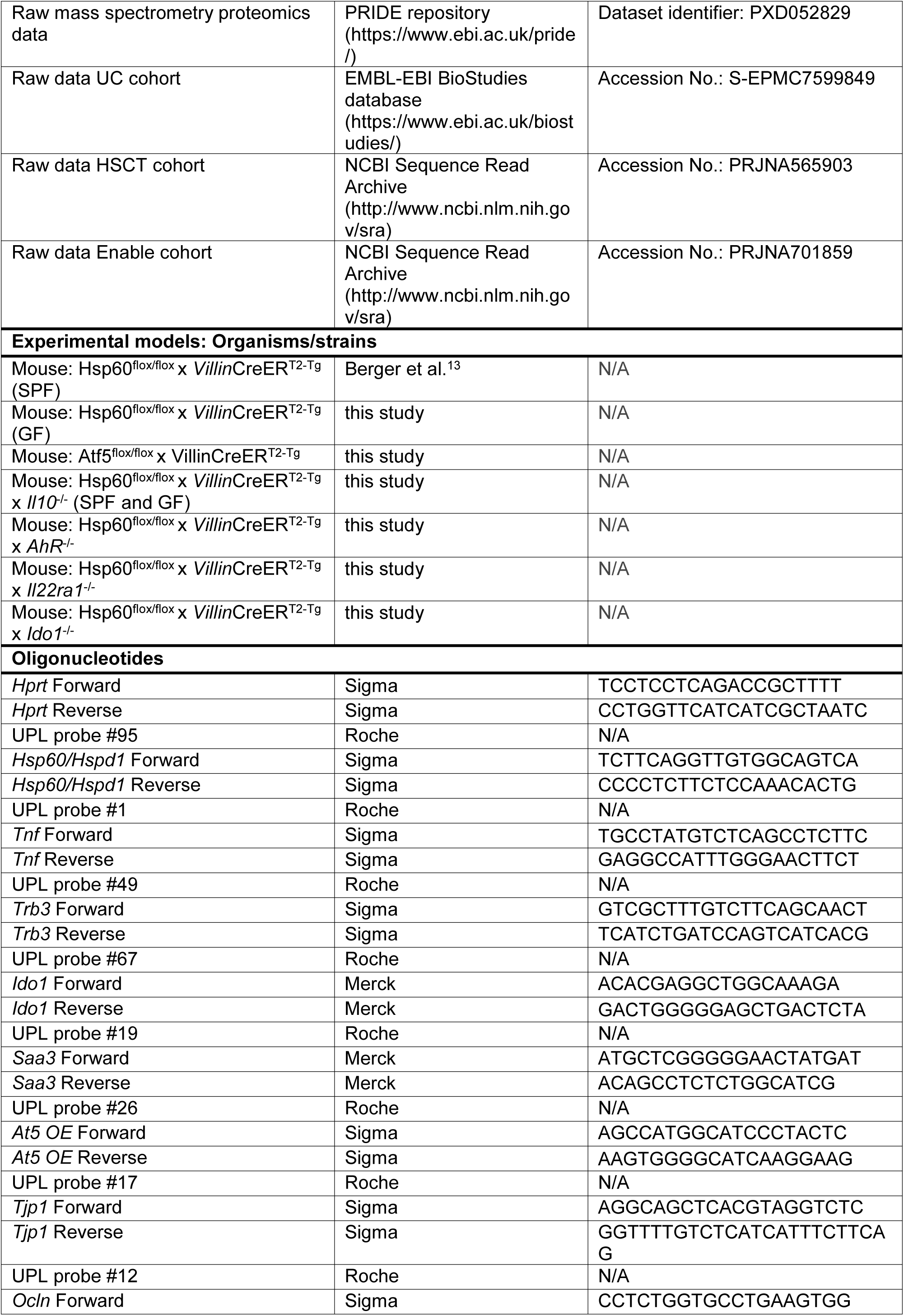
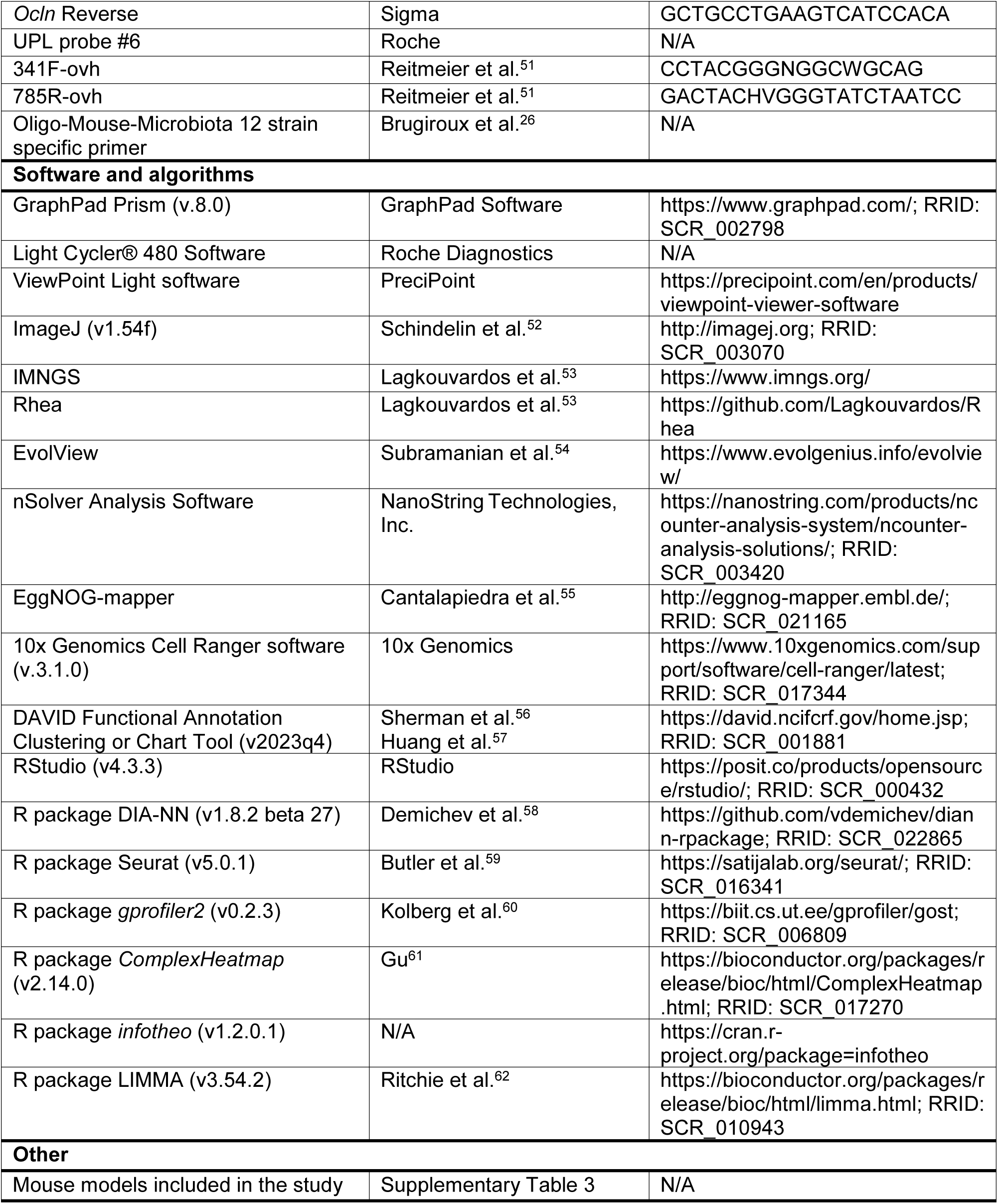

### RESOURCE AVAILABILITY

#### Lead contact

Further information and requests should be directed to and will be fulfilled by the lead contact Prof. Dr. Dirk Haller

#### Materials availability

All unique resources generated in this study may be available from the lead contact under a material transfer agreement.

#### Data and code availability

- The demultiplexed reads for the 16S rRNA amplicon sequencing dataset of the Biotherapy cohort have been deposited in the NCBI Sequence Read Archive and are publically available under the project number PRJNA701749. Raw data from HSCT and Enable cohorts have been deposited to the NCBI Sequence Read Archive [http://www.ncbi.nlm.nih.gov/sra] under the accession No. PRJNA565903 and No. PRJNA701859, respectively. Data from the UC cohort have been deposited to EMBL-EBI BioStudies database under the accession No. S-EPMC7599849. Software used to analyze the data are either freely or commercially available. Source code data are available from the corresponding author on request. Accession numbers are listed in the *key resources table*. Raw mass spectrometry proteomics data of bacterial metaproteomics have been deposited to the ProteomeXchange Consortium via the PRIDE partner repository (https://www.ebi.ac.uk/pride/) and are publicly available as of the date of publication. Accession numbers are listed in the *key resources table*. Single-cell RNA-sequencing raw data of mouse colonic tissue have been deposited at the EMBL-EBI ArrayExpress repository (https://www.ebi.ac.uk/biostudies/arrayexpress) and are publicly available as of the date of publication. Accession numbers are listed in the *key resources table*.
- This paper does not report original code. All used tools are cited in the respective methods and *key resources table*.
- Any additional information required to reanalyze the data reported in this paper is available from the lead contact upon request.

### EXPERIMENTAL MODEL AND STUDY PARTICIPANT DETAILS

#### Human cohorts

Microbial profiling of stool samples derived from 3 IBD patient cohorts were included: French Biotherapy cohort (N=147) and the two previously published Spanish HSCT cohort (N=153) ^25^, and German UC cohort (N=168) ^24^. The Enable cohort (PMID: 33195372 and PMID: 32619440), including healthy non-smoking individuals at different phases of life, recruited in Freising (Southern Germany) (N=817), was included for comparison. The Biotherapy cohort is a longitudinal cohort comprising a total of 80 patients with IBD (CD: N=41, UC: N=39) who were treated with biological therapy (CD patients: Infliximab N=9, Vedolizumab N=5, Ustekinumab N=12, Adalimumab N=13, NA=2; UC patients: Infliximab N=11, Vedolizumab N=10, Golimumab N=9, Adalimumab N=2, NA=7). Clinical disease monitoring and sampling were performed at three time points (weeks 0, 14, and 52 after treatment). This study was conducted by Prof. Matthieu Allez at the Hôpital Saint-Louis in Paris (NCT: 02693340). The transcriptome analysis of human biopsies from a multicenter prospective study (ClinicalTrials.gov [NCT03458195]) was coordinated by the REMIND group and previously published^29^. Microarray raw data are publicly available and accessed in NCBI’s Gene Expression Omnibus, through GEO series accession number GSE186582. All study methods were approved by the ethics committee of the hospital and complied with all relevant ethical regulations for studies with human research participants. Written informed consent was obtained from all subjects.

#### Ethics statement

All animal experiments, as well as maintenance and breeding of mouse lines, were approved by the Committee on Animal Health Care and Use of the state of Upper Bavaria (Regierung von Oberbayern; AZ ROB-55.2-2532.Vet_02-14-217, AZ ROB-55.2-2532.Vet_02-20-58, AZ ROB-55.2-2532.Vet_02-18-37) and performed in strict compliance with the EEC recommendations for the care and use of laboratory animals (European Communities Council Directive of November 24, 1986 (86/609/EEC)).

#### Animals and housing conditions

All mouse models are presented in Supplementary Table 3 and only male animals were used which were assigned to experimental group based on their respective genotypes. Briefly, Hsp60^flox/flox^ mice and Hsp60^flox/flox^ x *Villin*CreER^T2-Tg^ mice (to generate IEC-specific Hsp60 knockout mice via tamoxifen induction (Hsp60^Δ/ΔIEC^)) were generated as described previously^13^. In addition, Hsp60^Δ/ΔIEC^ mice were crossed with, respectively, whole body IL-10 knockout mice (*Il10^-/-^*), AhR knockout mice (*AhR^-/-^*), IL22ra1 knockout mice (*Il22ra1^-/-^*; kindly provided by Prof. S. Huber) and Ido1 knockout mice (*Ido1^-/-^*) to generate, respectively, Hsp60^Δ/ΔIEC^;Il10^-/-^, Hsp60^Δ/ΔIEC^;AhR^-/-^, Hsp60^Δ/ΔIEC^;Il22ra1^-/-^ and Hsp60^Δ/ΔIEC^;Ido1^-/-^ mice. To generate an IEC-specific and inducible overexpression of *Atf5*, Atf5^IEC^ (Atf5^flox/flox^ x *Villin*CreER^T2-Tg^), a vector construct with a CAG promoter and an N-terminal HA-Tag (allowing nuclear translocation and unhindered activity of Atf5 as a transcription factor) has been integrated into the Rosa 26 locus. Transgenic animals were crossed with *Villin*CreER^T2-Tg^. Inducible Cre-mediated recombination by tamoxifen feeding leads to the deletion of a promoter-associated STOP sequence, which prevents Atf5 expression. Hence, upon Cre-mediated recombination, Atf5 is expressed and translocated to the nucleus. All mice were housed under specific pathogen-free (SPF) or germ-free (GF) conditions according to the criteria of the Federation for Laboratory Animal Science Associations (FELASA) (12h light/dark cycles at 24–26°C) in the mouse facility at the Technical University of Munich (School of Life Sciences Weihenstephan). GF mice were housed in flexible film isolators ventilated via HEPA-filtered air at 22 ± 1°C. All mice received a standard diet (autoclaved V1124-300, Ssniff) *ad libitum*, autoclaved water and were sacrificed by CO_2_ or isoflurane. GF status was controlled by cultivating feces in Columbia Agar with 5% Sheep Blood (BD^TM^) before the start of the experiments and at sampling time points. All mice were housed with their respective littermates until 10 weeks of age. At the start of tamoxifen feeding and until sampling time point, all animals were single-housed.

#### Induction of postnatal recombination and monitoring of mice

For conditional Hsp60^flox/flox^ x *Villin*CreER^T2-Tg^, Hsp60^flox/flox^ x *Villin*CreER^T2-Tg^ x Il10^-/-^, Hsp60^flox/flox^ x *Villin*CreER^T2-Tg^ x AhR^-/-^, Hsp60^flox/flox^ x *Villin*CreER^T2-Tg^ x Il22ra1^-/-^, Hsp60^flox/flox^ x *Villin*CreER^T2-Tg^ x Ido1^-/-^ and Atf5^flox/flox^ x *Villin*CreER^T2-Tg^ mice and appropriate control mice were kept on phytoestrogen-reduced diet (V1154-300, Ssniff) for four weeks under SPF or GF conditions (from weeks 6 to 10 of age). Afterwards, 10 week-old male mice received 400mg tamoxifen citrate per kg chow feed (CreActive T400 (10mm, Rad), Genobios) *ad libitum* for 7 days. After the induction phase, tamoxifen diet was replaced with the phytoestrogen-reduced diet. During and after the induction phase, mice were monitored daily and aborted when a combined score considering weight loss, changes in stool consistency, general behavior, and general state of health was reached. Animals were sacrificed at the indicated time points.

#### Primary crypt isolation and intestinal organoid culture

Primary small intestinal crypts from mice (from 10- to 20-week old both male and female Hsp60^flox/flox^ x *Villin*CreER^T2-Tg^ mice and control mice) were isolated, and intestinal tissue (6-10cm) was inverted, digested with Ethylenediaminetetraacetic acid (EDTA, 2mM, 30min; 4°C), and mechanically dissociated. The supernatant was filtered via a 70μm cell strainer, pelleted by centrifugation, and embedded in Matrigel (BD Biosciences). 25µl Matrigel^®^-organoid suspension was plated in 48 well plates (Eppendorf) and incubated in IntestiCult™ Organoid Growth Medium (STEMCELL Technologies) in a humidified 5% CO_2_ atmosphere at 37°C. For subsequent passaging every 7-10 days, organoids were mechanically disrupted using a 1ml syringe with a 20G needle, pelleted by centrifugation, and embedded in fresh Matrigel. During experimental procedures, organoids were passaged three days before the start of the experiment and cultivated on self-made crypt culture medium (CCM). CCM contains advanced DMEM/F12 medium (Gibbco), 2mM GlutaMax (Gibbco), 10mM HEPES, penicillin, streptomycin and amphotericin (all Sigma-Aldrich) supplemented with N2 (1:100), B27 (1:50) (Gibbco), 1mM N-acetylcysteine (Sigma-Aldrich), 50ng/ml EGF (ImmunoTools), 100ng/ml noggin and 0.5µg/ml R-spondin 1 (both PeproTec).

#### Oligo-Mouse-Microbiota (OMM12) synthetic bacterial community

To prepare Gavages for the selective colonization of germ-free mice with the Oligo-Mouse-Microbiota (OMM^12^) synthetic bacterial community^27^, each bacteria was grown in Hungate tubes in Anaerobic Akkermansia Medium (AAM) ^26^ for 24-48h under anaerobic conditions at 37°C. The bacterial strains used are given in the *key resources table*. Bacterial growth was assessed using OD600, adjusted to mix the same amount of each bacteria needed with a density of at least 3×10^7^ bacteria/ml, and glycerol was added (20% v/v final concentration). The gavages were sealed with a rubber septum in N_2_-gased tubes and stored at -80°C until their transfer to the gnotobiotic facility.

### METHOD DETAILS

#### Selective colonizations of germ-free mice

For selective colonizations, GF male Hsp60^flox/flox^ x *Villin*CreER^T2-Tg^ mice and their appropriate control mice were rectally gavaged twice using 20 Gauge gavage needles (Fine Science Tools) at the age of 6 weeks with the Oligo-Mouse-Microbiota (OMM^12^) synthetic bacterial community^27^, *Limosilactobacillus reuteri* I49 (DSM 32035) or *Bacteroides caecimuris* I48 (DSM 26085) alone or in combination with *Akkermansia muciniphila* YL44 (DSM 26127). After the 2^nd^ gavage, induction of postnatal genomic recombination was performed at 10 weeks of age as described before. All bacterial gavages were prepared as described before for OMM^12^.

#### Tissue processing and histopathological analysis

Small intestinal and colonic tissues were removed immediately after euthanasia and dissected from adjacent tissue. The different parts of the gastrointestinal tract were cut open longitudinally, the content was removed, and the tissue processed as ‘swiss roles’. Tissues were fixed in 4% phosphate-buffered formalin for 48h, dehydrated, and embedded in melted paraffin (FFPE) or embedded in optical cutting temperature (OCT) medium without fixation. For histopathological assessment, 5μm sections were stained with hematoxylin (Mayer) and 0.2% eosin (ethanolic solution; both Medite) in an automated staining machine (Leica). Ileal and colonic sections were blindly assessed for inflammation (tissue pathology index) based on infiltration of mononuclear cells into the individual wall layers (*mucosa*, *submucosa* and *muscularis*), hyperplasia and atypia of the crypts, depletion of goblet cells and epithelial damage, resulting in a score from 0 (non-inflamed) to 12 (severely inflamed). In addition, ileal and colonic sections were blindly assessed for an aberrant epithelial architecture *via* crypt architectural distortion (crypt hyperplasia, shifted nucleus-plasma relations, enlarged nuclei, nucleoli, apical shifted mitoses) generating an aberrant epithelial index ranging from 0 (regular epithelium) to 4 (highly aberrant epithelium). Images were acquired by a PreciPoint M8 microscope (Precipoint).

#### Alcian blue / Periodicacid-Schiff (PAS-AB) staining

FFPE tissue sections (5μm) were stained with alcian blue (Fisher) for detection of acidic mucins (0.5% v/v in 3% acetic acid, pH=2.5, 5min) as well as periodic acid (0.5% v/v, 10min) co-stained with Schiff’s reagent (Sigma-Aldrich) for neutral mucins (15min). Nuclei were counterstained with hematoxylin. Images were acquired by a PreciPoint M8 microscope (Precipoint).

#### Staining procedures of FFPE sections of Hsp60, Ki67 and Ido1

For immunohistochemistry (IHC) and immunofluorescence (IF) stainings, FFPE tissue sections of intestinal ‘swiss roles’ (5µM) were deparaffinized and rehydrated in an automated staining machine (Leica). After heat-mediated antigen retrieval with 10mM citrate buffer (pH=6; 23min; 900 watt), sections were incubated for 10min with 3% H_2_O_2_ (Sigma-Aldrich) for IHC staining only. Subsequently, sections were blocked (blocking buffer consisting of 2.5ml 10x PBS, 1.25ml serum of the same species as the secondary antibody, 21.25ml dH_2_O, 75µl Triton X-100 (100%, Sigma-Aldrich)), and a primary antibody was applied overnight at 4°C. Antibody concentrations are given in the *key resources table* (antibody dilution buffer consisting of 4ml 10x PBS, 36ml dH_2_O, 0.4g BSA (Sigma-Aldrich), 120µl Triton X-100 (100%, Sigmal Aldrich)). After several washing steps, sections were incubated with the secondary antibody (fluorochrome- or horseradish peroxidase-coupled). In IF stainings, 4′,6-diamidino-2-phenylindole (DAPI; 1:1000; Sigma-Aldrich) was used for nuclei staining and added to the secondary antibody solution. IF stainings were visualized using the Fluoview FV10i microscope (Olympus) and images were processed using ImageJ (1.54f; http://imagej.org). For IHC stainings, antigen detection was performed by 3,3′-diaminobenzidine tetrahydrochloride (DAB)/metal concentrate (Fisher), and nuclei were counterstained with hematoxylin. IHC stainings were scanned and further analyzed using a PreciPoint M8 microscope (Precipoint).

#### Staining procedures of incorporated Bromodeoxyuridine (BrdU) on FFPE sections

BrdU solution (Invitrogen, Catalog#000103, 10µl/g of body weight) was administrated by intra-peritoneal injections 2h prior to sacrifice of all animals, in order to assess its incorporation into replicating IECs. FFPE tissue sections of intestinal ‘swiss roles’ (5µM) were deparaffinized and rehydrated in an automated staining machine (Leica). After heat-mediated antigen retrieval with 10mM citrate buffer (pH=6; 90°C on a heating plate for 10min), sections were cooled down for 20min and incubated for 10min with 3% H_2_O_2_ (Sigma-Aldrich). Using BD Pharmingen™ BrdU *In Situ* Detection Kit (AB_2868903). After washing, biotinylated anti-BrdU antibody was added for 1h at room temperature in a humidified chamber. After 3 washes in PBS, the secondary Streptavidin-HRP was incubated for 30min at room temperature in a humidified chamber and antigen detection was performed with DAB reagent from the kit. Nuclei were counterstained with hematoxylin. Stainings were scanned and further analyzed using a PreciPoint M8 microscope (Precipoint).

#### Staining procedure of internucleosomal DNA fragmentation

*In situ* apoptosis of IECs was assessed using Apo-BrdU *In Situ* DNA Fragmentation Assay Kit (BioVision, Catalog #K401-60). FFPE tissue sections of intestinal swiss roles (5µM) were deparaffinized and rehydrated in 0.85% NaCl for 5min. After incubation in 4% paraformaldehyde for 15min, slides were washed and processed as described by the manufacturer’s instructions. DAPI was used to co-stain nuclei. Stainings were visualized using the Fluoview FV10i microscope (Olympus) and images were processed using ImageJ (1.54f; http://imagej.org).

#### Staining procedures of tight junctional proteins

Colonic cryosections (6µm) were fixed for 30 min with 4% paraformaldehyde at room temperature, permeabilized for 30 min in 0.1% Triton X-100 at 4 °C and blocked for 1h at room temperature (blocking buffer consisting of 2.5ml 10x PBS, 1.25ml serum of the same species as the secondary antibody, 21.25ml dH_2_O, 75µl Triton X-100 (100%, Sigma-Aldrich)). Subsequently the primary antibody was applied overnight at 4°C (antibody dilution buffer consisting of 4ml 10x PBS, 36ml dH_2_O, 0.4g BSA (Sigma-Aldrich), 120µl Triton X-100 (100%, Sigmal Aldrich)). Antibody dilutions are given in the *key resources table*. After several washing steps, and 30min blocking at room temperature sections were incubated with the secondary antibody (fluorochrome-coupled). DAPI (1:1000; Sigma-Aldrich) was used for nuclei staining and added to the secondary antibody solution. After post-fixation (4% paraformaldehyde at room temperature), slides were mounted with Aquatex^®^ (Sigma-Aldrich). IF stainings were visualized using the Fluoview FV10i microscope (Olympus) and images were processed using ImageJ (1.54f; http://imagej.org).

#### mRNA isolation and quantitative real-time PCR

mRNA of total colonic tissue or small intestinal organoids was isolated using the NucleoSpin RNAII kit (Macherey-Nagel) according to the manufacturer’s instructions. Quantitative Real-time PCR (qRT–PCR) was performed in duplicates using 1μl cDNA on a Light Cycler 480 system (Roche Diagnostics), applying the Universal Probe Library system according to the manufacturer’s instructions. Primer sequences and probes are provided in the *key resources table*. Relative induction of gene mRNA expression was calculated by the 2^-ΔΔCt^ method using the expression of the housekeeper *Hprt* for normalization.

#### Quantitative qPCR of bacterial 16S rRNA genes

Quantitative PCR of the twelve different bacteria from the OMM^12^ consortium was performed using 10ng of gDNA as a template. Strain-specific 16S rRNA primers and hydrolysis probes were used for amplification^26^. For duplex qPCR assays, hydrolysis probes were 5′-labelled with either 6-carboxyfluorescein (FAM) or 6-carboxyhexafluorescein (HEX). qPCR standard curves were determined using linearized plasmids as DNA templates and run on a Light Cycler 480 system (Roche Diagnostics) in duplicates. A standard curve was then used for absolute quantification of 16S rRNA gene copy numbers of individual strains. *Acutalibacter muris* KB18 (DSM 26090) signals were below detection limits.

#### Mouse ELISA SAA

Saa concentration in murine plasma was quantified using a commercially available ELISA kit (ICL Inc.; Cat. num. E-90SAA) according to the manufacturer’s instructions.

#### Metagenomic DNA extraction from colonic content

DNA was extracted from frozen mouse colonic content by bead-beating followed by a modified version of the protocol by Godon *et al*., as previously described^10^. Briefly, a volume of 600µl DNA stabilizing solution (Stratec Biomedical) was added to the colonic content aliquots in 2ml screw-cap polypropylene microcentrifuge tubes containing 500mg sterile Silica beads (0.1-mm-diameter; BioSpec Products) on ice. For absolute analysis, spikes were added to the samples (1.5 ng/µl). After the addition of 250µl of 4M guanidine thiocyanate – 0.1M Tris (pH=7.5) and 500µl of 5% *N*-lauroyl sarcosine – 0.1M phosphate buffer (pH=8.0), colonic content suspensions were vortexed and incubated at 70°C for 1h with constant shaking. The mixture was mechanically disrupted by bead beating using a FastPrep®-24 bead beater (MP Biomedicals) supplied with a 24 × 2ml cooling adapter three times each for 40s at a speed of 6.5m/s. 15mg polyvinylpolypyrrolidone (PVPP, Sigma-Aldrich) was added as polyphenol adsorbent, and the suspension was centrifuged for 3min at 15,000*g* at 4°C. The supernatant was recovered and centrifuged for 3min at 15,000*g* at 4°C. To remove bacterial RNA, a volume of 2µl RNAse (10mg/ml) was added to 500 µl clear supernatant and incubated at 37°C for 30min with constant shaking. Finally, the genomic DNA was purified using the NucleoSpin® gDNA clean-up kit (Macherey Nagel) following the manufacturer’s instructions. The concentration and purity of the extracted DNA were determined using the NanoDrop® Spectrophotometer ND-1000 (ThermoFisher Scientific).

#### 16S#rRNA gene amplicon sequencing

After genomic DNA extraction, amplicon libraries (V3/V4 region) were amplified by PCR (25 cycles), purified using the AMPure XP system (Beckmann), pooled in an equimolar amount, and sequenced in paired-end mode (PE275) using a MiSeq system (Illumina, Inc.) according to the manufacturer’s instructions and 25% (v/v) PhiX standard library. Raw sequences were processed using IMNGS based on the UPARSE approach^63^. Sequences were demultiplexed, trimmed to the first base with a quality score <3 and then paired. Sequences with less than 300 and more than 600 nucleotides and paired reads with an expected error >3 were excluded from the analysis. The remaining reads were trimmed by 5 nucleotides on each end to avoid GC bias and nonrandom base composition. The presence of chimeras was tested using UCHIME^64^. Clustering of zero-radius operational taxonomic units (zOTUs) was done at 97% sequence similarity. Taxonomic binning was assigned at an 80% confidence level using the RDP classifier^65^ and compared to the SILVA ribosomal RNA gene database project^66^. Analysis was performed using the R-package Rhea^53^. In brief, rarefaction curves were used to assess sequencing depth and eliminate low-quality reads. zOTU counts were normalized, and percentage relative abundance was computed. Alternatively, absolute abundance was computed from non-normalized zOTUs counts. Beta-diversity analysis was used to determine the diversity between groups based on generalized UniFrac distances. Diversity within species was calculated based on species richness. EvolView was used to visualize phylogenetic trees^54^.

#### Shallow shotgun metagenomics

Shallow shotgun metagenomic sequencing was performed on a NovaSeq machine (1G raw data with paired-end 150bp) using gDNA isolated from murine colonic content. After demultiplexing the data, adapters were trimmed from the raw sequencing reads using Trim Galore (https://github.com/FelixKrueger/TrimGalore). Reads were subsequently processed with the KneadData quality control pipeline to remove low-quality read bases and murine contamination (https://bitbucket.org/biobakery/kneaddata/wiki/Home). Taxonomic profiling was performed using the MetaPhlAn3.0.13 classifier^67^, which unambiguously classifies metagenomic reads to taxonomies based on a database of clade-specific marker genes derived from 17,000 microbial genomes (∼13,500 bacterial and archaeal, ∼3,500 viral, and ∼110 eukaryotic). Species-level assignments were extracted, and data was centered log ratio scaled.

#### RNA Microarray analysis

Total RNA was prepared from isolated colonic epithelial cells of Hsp60^Δ/ΔIEC^ and Hsp60^fl/fl^ mice on day 2 with the column-based NucleoSpin RNAII kit (Macherey-Nagel) according to the manufacturer’s instructions. RNA concentration and quality were determined by spectrophotometric analysis (ND-1000, PEQLAB). RNA integrity number (RIN) was >8 for all samples (Agilent 1200 bioanalyzer, Alpha Metrix Biotech). Labeling of total RNA (100ng) was performed with the Ambion WT Expression kit (Life Technologies), and samples were hybridized to Affymetrix GeneChip Mouse Gene 1.1 ST arrays (Affymetrix) and scanned on an Affymetrix GeneTitan instrument. Probe sets were defined according to Dai *et al.* ^68^. Probes were annotated and assigned to unique gene identifiers (Entrez IDs). The probes present on the Mouse Gene 1.1 ST arrays represent 21,225 Entrez IDs. Arrays were normalized with the Robust Multi-array Average (RMA) method. Only probe sets with normalized signal intensities above 20 on at least five arrays, interquartile range > 0.2, and at least 5 probes per probe set were selected for further analysis.

#### Generation of murine cecal water

To generate murine cecal water, approximately 150mg frozen cecal content was mixed with 3 sterile glass beads (2mm diameter; Glasschrott) and sterile PBS (10% w/v), followed by vigorous vortexing for 1min. To remove food debris, the samples were centrifuged (5min, 500g, 4°C), followed by the transfer of the supernatant into new tubes. After a second centrifugation step (5min, 8000g, 4°C), bacteria were pelleted, and the supernatant was distinguished as cecal water.

#### Induction of *ex vivo* recombination and stimulation of organoids

*Ex vivo* induction of the *Hsp60* deletion in IECs was achieved by adding 500nM (Z)-4-hydroxytamoxifen (4-OHT; LKT) to the culture medium. After 24h, organoid culture media was changed to fresh CCM. Hsp60 is completely gone on protein level after 4 days. Five days after *ex vivo Hsp60* deletion, organoids were treated for 24h with Tnf (50ng/ml; PeproTech Inc.), Ifnγ (1ng/ml; PeproTech Inc.), lipopolysaccharides (LPS; 1µg/ml; Sigma-Aldrich) or murine cecal water (dilution 1:500) and subsequently harvested for RNA isolation.

#### Measurement of living cells and cellular ATP content

Live- and dead-cell protease activity was measured using the MultiTox-Fluor Cytotoxicity Assay, and ATP content was measured using the CellTiter-Glo Luminescent Cell Viability Assay (both from Promega), according to the manufacturer’s instructions in a 48-well format. Fluorescence and luminescence were measured using the Tecan infinite M200 (Tecan Group Ltd.) and the i-control™ Microplate Reader software (Tecan).

#### Colonic content protein extraction and digestion for metaproteomics analysis

Protein extraction from the mouse colonic content was performed as previously described^69^. Briefly, 30mg of each sample was taken (in the case of less than 30mg of available material, the whole sample was used). The samples were mixed with 200μl of lysis buffer (5% SDS, 2M Urea, 50mM Tris-HCl, Protease Inhibitor 1x) and vortexed vigorously for 1min. Samples were then placed in a Thermomixer (Eppendorf, Hamburg, Germany) for 15min at 70°C with 1,200 rpm agitation. When finished, samples were ultrasonicated using a Bioruptor Pico (Diagenode, Seraing, Belgium; program: 15 cycles of 30 seconds “ON” and 30 seconds “OFF”, frequency level: Low, water temperature: +20°C). Samples were centrifuged at 16,000g for 5min at room temperature. Supernatants were transferred into new tubes and submitted to a new round of centrifugation in the same conditions. Collected supernatants were stored at -80°C until further use.

For protein clean-up and digestion, a modified version of the single-pot, solid-phase-enhanced sample preparation (SP3) method described by Hughes et *al.* ^70^ was used. Briefly, 50µg protein was taken into a 1.5ml LoBind tube (Eppendorf, Hamburg, Germany), and the sample volume was added up to 90µl with lysis buffer. The colonic content protein sample was subjected to protein reduction (5mM Dithiothreitol, DTT, 30min incubation at 60°C) followed by alkylation (20mM Iodoacetamide, IAA, 30min at room temperature in the dark). The remaining IAA in the sample was quenched with the addition of DTT to a final concentration of 5mM. 10µl of pre-mixed Sera-Mag SpeedBead beads (50mg/ml, Cytiva, Marlborough, Massachusetts) were added to 50µg protein sample. To initiate the binding of proteins to the beads, 100µl absolute ethanol was added immediately, followed by incubation in the Thermomixer at 24°C for 5min with 1,000rpm agitation. The supernatant was removed after 2min resting on a magnetic rack, and the beads were rinsed three times with 50μl of 80% ethanol. Rinsed beads were reconstituted in 50μ digestion buffer (50mM ammonium bicarbonate, pH 8). Protein digestion was performed with 2μg of sequencing-grade trypsin/LysC (Promega, Madison, USA) for 18 hours at 37°C with 950rpm agitation. After digestion, ACN was added to each sample to a final concentration of 95%. Mixtures were incubated for 8min at room temperature and then placed on a magnetic rack for 2min. The supernatant was discarded, and the beads were rinsed with 900μL of 100% ACN. Rinsed beads were reconstituted in 20μl LC-MS grade water to elute the peptides. The peptide concentration was measured in duplicate using NanoPhotometer N60 (Serial number: TG2022, Implen, Munich, Germany) at 205nm. Peptide samples were acidified with formic acid (FA) to a final concentration of 0.1% and stored at -20°C until LC-MS/MS analysis.

#### Liquid Chromatography Tandem Mass Spectrometry (LC-MS/MS) setup

Nanoflow reversed-phase liquid chromatography (Nano-RPLC) was performed on a NanoElute system (Bruker Daltonik, Bremen, Germany). 250ng of digested peptides were separated with a 60min gradient on a 25cm x 75µm column packed with 1.6µm C18 particles (IonOpticks, Fitzroy, Australia). Mobile solvent A consisted of 100% water (LC-MS grade), 0.1% FA and mobile phase B of 100% acetonitrile, 0.1% FA. The flow rate was set to 250nl/min or the whole separation apart from the last 7min where the flow was set to 400nl/min. The mobile phase B was linearly increased from 4 to 20% in the first 35min, followed by another linear increase to 35% within 17min and a steep increase to 85% in 0.5min. Then a flow rate of 400nl/min with 85% of mobile solvent B was maintained for 7min to elute all hydrophobic peptides. NanoElute LC was coupled with a hybrid TIMS quadrupole TOF mass spectrometer (timsTOF Pro, Bruker Daltonik, Bremen, Germany) via a CaptiveSpray ion source. Samples were analyzed in data-independent acquisition (DIA) modes coupled with parallel accumulation serial fragmentation (PASEF). For the acquisition, the TIMS analyzer was operated in a 100% duty cycle with equal accumulation and ramp times of 100ms each. The ion mobility range was set to 0.6 - 1.4 (1/k0). Precursors with m/z between 3500 and 1100 were defined in 13 scans containing 30 isolation steps with an isolation window of 25 Th in each step. The acquisition time of each DIA-PASEF scan was set to 100ms, which led to a total cycle time of around 1.5 sec. The collision energy was ramped linearly from 59 eV at 1/k0 = 1.6 to 20 eV at 1/k0 = 0.6.

#### Epithelial cell isolation from colonic mouse tissue for scRNA-seq library preparation and sequencing

The entire colon was removed from either Hsp60^Δ/ΔIEC^ or Hsp60^fl/fl^ mice, weighed and stored in Dulbecco’s Modified Eagle Medium (DMEM). After being washed with PBS and minced on ice, tissue specimens were dissociated and enzymatically digested (Multi Tissue Dissociation Kit using gentleMACS™ Dissociator, Miltenyi Biotec). After centrifugation and resuspension in PBS+2% FCS, the cell suspension was strained through a 70-µm strainer, spun down and resuspended in 2% FCS/PBS including RNase inhibitor (New England Biolabs, catalog no. M0314L,1:100). Debris removal solution (Miltenyi Biotec, catalog no. 130-109-398) was used to remove cell debris from the dissociated tissue. Then the dead cell removal kit (Miltenyi, catalog no. 130-090-101) was used to enrich live cells. CD45+ cells were sorted out using the CD45 Microbeads kit (Miltenyi Biotec, catalog no. 130-052-301). Remaining cells were counted and diluted in 2% FCS/PBS and up to 30,000 cells were loaded per lane on a 10× Chromium chip to generate gel beads in emulsion (GEMs). Single-cell GEM generation, barcoding and library construction were performed by using 10× Chromium Single Cell 3′ v.3 chemistry according to the manufacturer’s instructions. Complementary DNA and generated libraries were checked for sample size and quality on an Agilent Bioanalyzer 2100 using HS DNA Kit (Agilent). Libraries were sequenced on Illumina NovaSeq 6000 S2 (Paired ends (PE), 28 + 94 bp).

### QUANTIFICATION AND STATISTICAL ANALYSIS

#### Quantification of Hsp60 expression

Hsp60 protein expression was quantified in Hsp60 IHC stainings of colonic tissue sections using the ViewPoint Light software. To categorize Hsp60 expression in crypts, fifty well-oriented crypts were categorized as positive, half-positive, or negative for Hsp60 expression. Half-positive crypts were crypts with a positive signal in only the left or right side of the crypt.

#### Quantification of goblet cells

The number of goblet cells was counted based on PAS-AB staining. Fifty well-oriented crypts were selected, and PAS-positive cells were counted. The number of PAS-positive cells was calculated as a total number per 100μm^2^.

#### Quantification of crypt depth

Crypt depth was analyzed in H&E stainings of colonic tissue sections. In fifty well-oriented crypts, the length from the crypt base to crypt top was measured using the ViewPoint Light software, and the average length was calculated for each section.

#### Quantification of Ki67-positive and BrdU-positive cells

To assess proliferation rate, Ki67-positive and BrdU-positive IEC were quantified based on, respectively, Ki67 and BrdU stainings of colonic tissue sections using the ViewPoint Light software. To determine Ki67- or BrdU-positive crypts, fifty well-oriented crypts (25 crypts in proximal and 25 in distal colon) were categorized as positive or negative for Ki67 or BrdU expression, and the number of Ki67-positive or BrdU-positive cells was calculated per crypt.

#### NanoString^®^ analysis

Multiplexed gene expression of a panel of 768 metabolic genes was analyzed using the nCounter® Metabolic Pathway Panel (NanoString Technologies, Inc.) according to the manufacturer’s instructions. In brief, 125ng isolated RNA of total colonic tissue underwent code set hybridization and was processed using a nCounter® MAX/FLEX system (NanoString Technologies, Inc.). Raw data were subjected to quality control, followed by normalization and analysis applying the nSolver Analysis Software (NanoString Technologies, Inc.).

#### DIA-PASEF data processing

DIA-NN^58^ (version 1.8.2 beta 27) was used to process DIA-PASEF data in library-free mode against 3 reference proteomes (downloaded from Uniprot on 20240105): Mus musculus (UP000000589), *Akkermansia muciniphila* (UP000001031), and *Bacteroides caecimuris* (UP000092631). A deep learning-based method was used to predict theoretical peptide spectra along with their retention time and ion mobility. Trypsin/P was used for *in silico* digestion with an allowance of a maximum of 1 missed cleavages. Variable modifications on peptides were set to methionine oxidation and N-term acetylation, while N-term methionine excision and carbamidomethylation on cysteine were fixed modifications. The maximum number of variable modifications on a peptide was set to 2. Peptide length for the search ranged from 7 to 30 amino acids. Aligned with the DIA-PASEF acquisition method, m/z ranges were specified as 350 to 1100 for precursors and 100 to 1700 for fragment ions. Both MS1 and MS2 mass accuracy were set to automatic determination. Protein inference was set to “Protein names (from FASTA)” and the option of “Heuristic protein inference” was unchecked. Match-between-run (MBR) was checked for cross-run analysis. RT-dependent cross-run normalization and QuantUMS (high precision) options were selected for quantification.

The DIA-NN search output was further processed with the R package, DIA-NN (https://github.com/vdemichev/diann-rpackage), to extract the quantitative intensity or for all identified peptides and protein groups with q-value ≤ 0.01 as criteria at precursor and protein group levels.

#### Functional annotation of 2 bacterial proteomes

The aforementioned proteome fasta files of *Akkermansia muciniphila* and *Bacteroides caecimuris* were imported to EggNOG-mapper^55^ (http://eggnog-mapper.embl.de/) for orthologue functional annotation with default settings. The functions of annotated proteins were then mapped to the DIA-PASEF identifications.

#### scRNA-seq data preprocessing, quality control and analysis

Alignment of the scRNA-seq data to the mouse reference genome (mm10, release 108.20200622), filtering, barcode and unique molecular identifier counting was performed using the 10x Genomics Cell Ranger software (v.3.1.0). All further analysis were performed in R v4.3.3 using Seurat v5.0.1^59^. Briefly, data generated from 10× pipeline were read and features which were detected in less than 3 cells were removed. Cells with less than 200 features detected were removed. RNA data was normalized with the Seurat method *sctransform* v2^71^. A threshold of more than 30% mitochondrial DNA was used to filter damaged cells. Additionally, doublets were removed with *scDblFinder* v1.16.0^72^ as well as cells with more than 20000 reads. Samples were integrated using Harmony v1.2.0^73^. Clusters were identified using the Louvain algorithm^74^ implemented in the *FindClusters()* Seurat method with a resolution of 0.1, based on a shared nearest neighbor graph (determined by the *FindNeighbors()* method). Subsequent sub-clustering of larger clusters using the above-explained approach was performed. Differentially expressed genes per cluster and condition were identified using the *FindMarkers()* Seurat method with MAST v1.27.1^75^. For the functional annotation of clusters, all differentially expressed genes (DEGs) with an adjusted p_value<0.01 and log_FC>1.5 were selected and annotated using DAVID Functional Annotation Clustering or Chart Tool, v2023q4 (https://david.ncifcrf.gov/home.jsp). Pseudobulk analysis of target genes was performed using the *AggregateExpression()*Seurat function.

#### Mapping of the murine mucosal signature to human Crohn’s disease samples

To study the expression of the murine signature in human Crohn’s disease patients, RMA normalized, log2 transformed microarray data were obtained from Ngollo *et al*.^29^. Samples from two different locations at the time of surgery were considered: the inflamed ileum [M0I] (n=196) and the ileal margin [M0M] (n=147). The latter served as control samples of uninflamed tissue. The signature genes were mapped from mouse to human gene names with the *gprofiler2* [version 0.2.3] R package ^60^. All but three of the 35 signature genes were successfully mapped to human orthologs in the microarray data. The three exceptions were *H2-Q10* and *Scd1*, for which no orthologs could be found, and *Slc16a3*, whose ortholog was not detected in the human microarray data. To visualize the expression of the signature’s human orthologs in a heatmap, the expression values were gene-wise scaled. Subsequently, the mean expression value of each location (M0I and M0M) was calculated. The *ComplexHeatmap* [version 2.14.0] R package ^61^ was used to generate the heatmap and to perform hierarchical clustering of the genes based on their Euclidian distance and complete linkage. The rows were annotated based on the expression patterns observed between the Hsp60^Δ/ΔIEC^ and Hsp60^fl/fl^ mice. To assess the significance of the number of genes with a concordant expression pattern in mice and humans, mutual information was calculated with the *infotheo* [version 1.2.0.1] R package (https://cran.r-project.org/package=infotheo). An empirical p-value for the mutual information was obtained through 10,000 Monte-Carlo permutations. For the differentially expressed genes (DEGs) analysis between M0I and M0M groups, the LIMMA [version 3.54.2] R package was used based on paired moderated t-statistics with Benjamini−Hochberg FDR correction for multiple testing ^62^. The DEGs were selected as biologically significant using adjusted *P* < 0.05. The code used for the analysis of the mapping of the murine gene signature to human samples has been deposited at GitHub repository (https://github.com/biomedbigdata/gene_signatures) and is publicly available.

#### Statistical analysis

Statistical analysis was performed with GraphPad Prism (version 8.0). Statistical tests used were: unpaired t-test, analysis of variance (ANOVA) followed by pairwise comparison testing (Tukey), or by Kruskal-Wallis test followed by Dunn’s multiple comparison. *P* < 0.05 was considered significant. \**P* < 0.05; \*\**P* < 0.01; \*\*\**P* < 0.001; \*\*\*\**P* < 0.0001. Data are presented as mean ± s.d., or in the case of weight curves, as mean ± sem.

---

**Figure S1:**
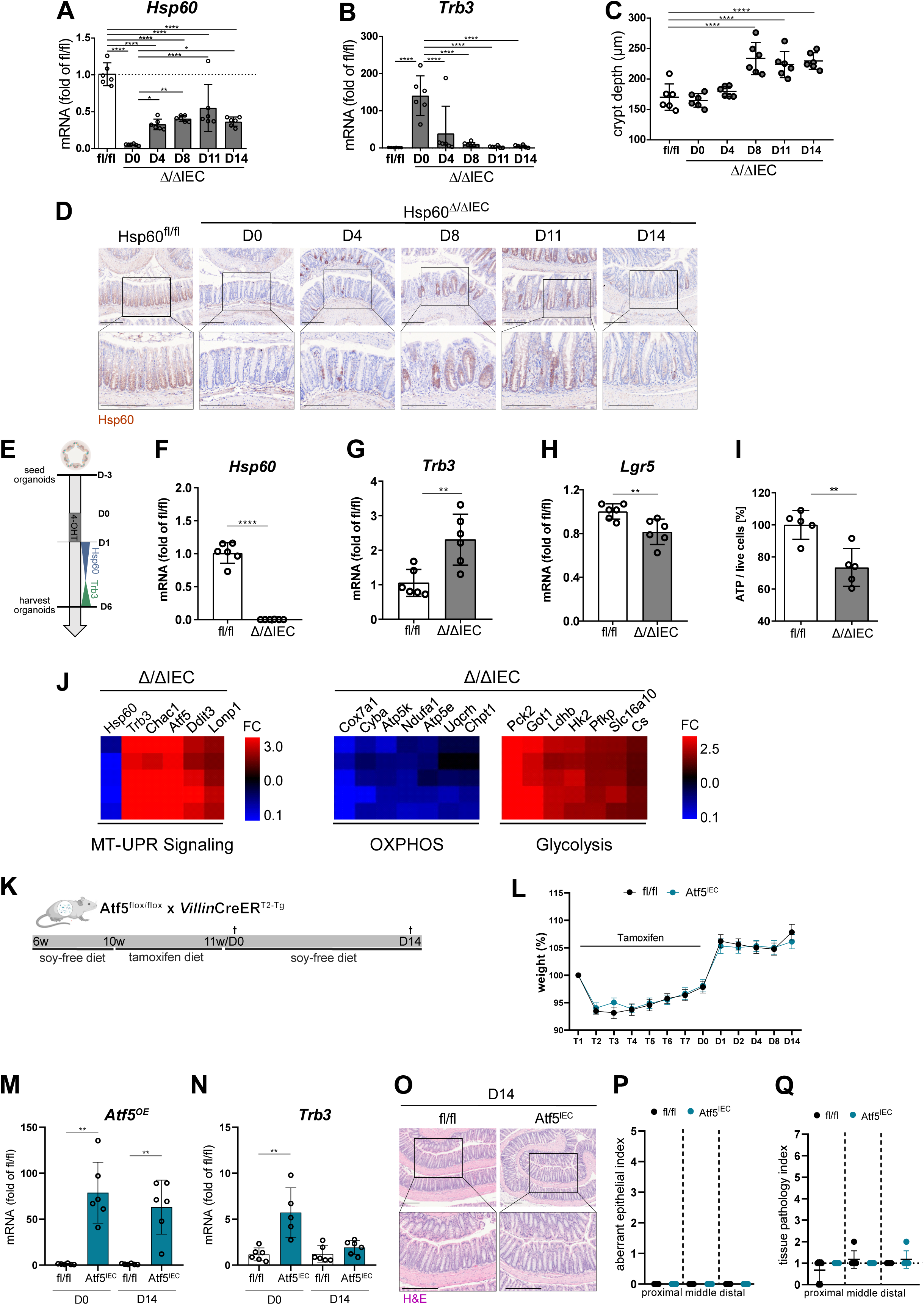
Hsp60 loss drives changes in cellular metabolism but mild MT-UPR induction does not lead to tissue injury. Related to Figure 1. **A, B,** qPCR of (**A**) the mitochondrial chaperone *Hsp60* and (**B**) the MT-UPR marker gene *Trb3* using mRNA of whole colonic tissue from day 0 to 14 in Hsp60^Δ/ΔIEC^ and Hsp60^fl/fl^ mice. **C,** Quantification of crypt depth of H&E stainings of colonic tissue of Hsp60^Δ/ΔIEC^ and Hsp60^fl/fl^ mice from day 0 to 14. **D,** Representative images of immunohistochemistry staining of Hsp60 in colonic tissue of Hsp60^Δ/ΔIEC^ and Hsp60^fl/fl^ mice from day 0 to 14. Brown color shows specific Hsp60 signal, nuclei are stained with hematoxylin. Scale bars: 200µm. **E,** Experimental setup. Small intestinal crypts were isolated and seeded in Matrigel^®^, followed by Hsp60 deletion *ex vivo* via tamoxifen addition to the media. Organoids were analyzed 6 days after *ex vivo* Hsp60 deletion. **F, G, H,** qPCR of (**F**) the mitochondrial chaperone *Hsp60*, (**G**) the MT-UPR marker gene *Trb3* and (**H**) the stem cell marker *Lgr5*. **I,** ATP measured per live cells at day 6 in the respective genotypes. **J,** Colonic crypts of Hsp60^Δ/ΔIEC^ and Hsp60^fl/fl^ mice were isolated at day 2, and RNA microarray analysis of genes related to MT-UPR signaling or OXPHOS and glycolysis was performed (data presented as fold change of Hsp60^Δ/ΔIEC^ over Hsp60^fl/fl^). **K,** Experimental setup. Atf5^flox/flox^ x *Villin*CreER^T2-Tg^ mice (Atf5^IEC^; Ctrl: Atf5^fl/fl^) were fed a soy-free diet for 4 weeks, followed by a tamoxifen diet at the age of 10 weeks for 7 days to induce Atf5 overexpression in IECs. Mice were sampled 0 and 14 days after tamoxifen feeding. **L,** Relative weight curve of Atf5^IEC^ and Atf5^fl/fl^ mice from the start of tamoxifen feeding until sampling time point. **M, N,** qPCR of (**M**) the mutant mRNA of the overexpressed transcription factor *Atf5* and (**N**) the MT-UPR marker gene *Trb3* using mRNA of whole colonic tissue from day 0 and 14 in Atf5^IEC^ and Atf5^fl/fl^ mice. **O,** Representative images of H&E staining of colonic tissue of Atf5^IEC^ and Atf5^fl/fl^ mice at day 14. Scale bars: 200µm. **P, Q,** H&E staining of colonic tissue of Atf5^IEC^ and Atf5^fl/fl^ mice on day 14 were histologically scored for (**P**) an aberrant epithelium and (**Q**) tissue inflammation. Data are represented by (**A-C, M, N, P, Q**) mean ± s.d. or (**L**) mean ± sem of six or **(J**) four biological replicates and (**F-I**) six technical replicates. (**A-C, P, Q**). P-values were calculated by ANOVA followed by pairwise comparison testing (Tukey). (**M, N**) P-values were calculated by Mann-Whitney test. *P<0.05, **P<0.01, ***P<0.001, ****P<0.0001. D: day; Hsp60: heat shock protein 60; 4-OHT: tamoxifen; IECs: intestinal epithelial cells; MT-UPR: mitochondrial unfolded protein response; Trb3: tribbles pseudokinase 3; Lgr5: leucine-rich repeat-containing G-protein coupled receptor 5; ATP: Adenosine triphosphate; OXPHOS: oxidative phosphorylation; Atf5: Activating Transcription Factor 5; Atf5^OE^: Atf5 overexpression transgene mRNA; H&E: hematoxylin and eosin.

**Figure S2:**
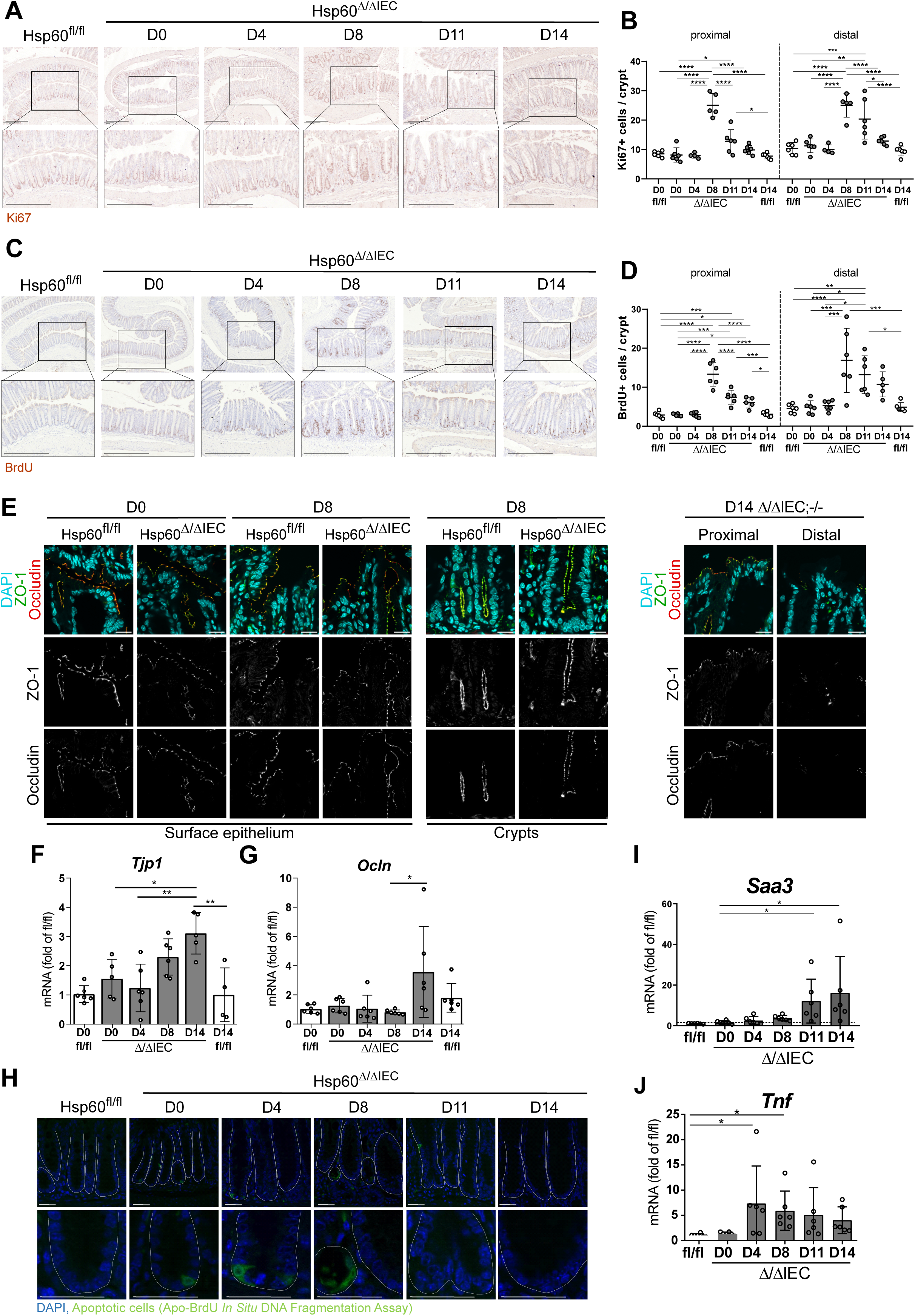
Barrier integrity is maintained during metabolic injury. Related to Figure 1. **A, C,** Representative images of immunohistochemistry staining of the proliferation markers (**A**) Ki67 and (**C**) incorporated BrdU in distal colonic tissue of Hsp60^Δ/ΔIEC^ and Hsp60^fl/fl^ mice from day 0 to 14. Brown color shows specific Ki67 or BrdU signal, nuclei are stained with hematoxylin. Scale bars: 200µm. **B, D**, Quantification of (**B**) Ki67-positive cells per crypt and (**D**) incorporated BrdU-positive cells per crypt in colonic tissue of Hsp60^Δ/ΔIEC^ and Hsp60^fl/fl^ mice from day 0 to 14. **E,** Representative immunofluorescent co-staining of tight junction proteins ZO-1 (green) and Occludin (red) as well as DAPI (blue) (left panel) at day 0 and 8 at the epithelium surface and in the bottom of crypts in distal colonic tissue of Hsp60^Δ/ΔIEC^ and Hsp60^fl/fl^ mice and (right panel) at day 14 in proximal and distal colonic tissue of Hsp60^Δ/ΔIEC^;Il10^- /-^ mice. Scale bars: 20µm. **F, G,** qPCR of tight junctional protein genes (**F**) *Tjp1* and (**G**) *Ocln* using mRNA of whole colonic tissue from day 0 to 14 in Hsp60^Δ/ΔIEC^ and Hsp60^fl/fl^ mice. **H,** Detection of apoptosis by Apo-BrdU *In Situ* DNA Fragmentation Assay (in green) on colonic tissue of Hsp60^Δ/ΔIEC^ and Hsp60^fl/fl^ mice from day 0 to 14. DAPI: blue. **I, J,** qPCR of the inflammatory marker genes (**I**) *Saa3* and (**J**) *Tnf* using mRNA of whole colonic tissue from day 0 to 14 of Hsp60^Δ/ΔIEC^ and Hsp60^fl/fl^ mice. Data are represented by mean ± s.d. of six biological replicates. (**B, D, F**) P-values were calculated by ANOVA followed by pairwise comparison testing (Tukey). **(G, I, J**) P-values were calculated by Kruskal-Wallis followed by pairwise comparison testing (Dunn). *P<0.05, **P<0.01, ***P<0.001, ****P<0.0001. D: day; Hsp60: heat shock protein 60; IECs: intestinal epithelial cells; BrdU: Bromodeoxyuridine; ZO-1: Zonula Occludens-1; Tjp1: tight junction protein1; Ocln: Occludin; DAPI: 4′,6-diamidino-2-phenylindole; *Saa*: serum amyloid A; *Tnf*: tumor necrosis factor; IL10: interleukin-10.

**Figure S3:**
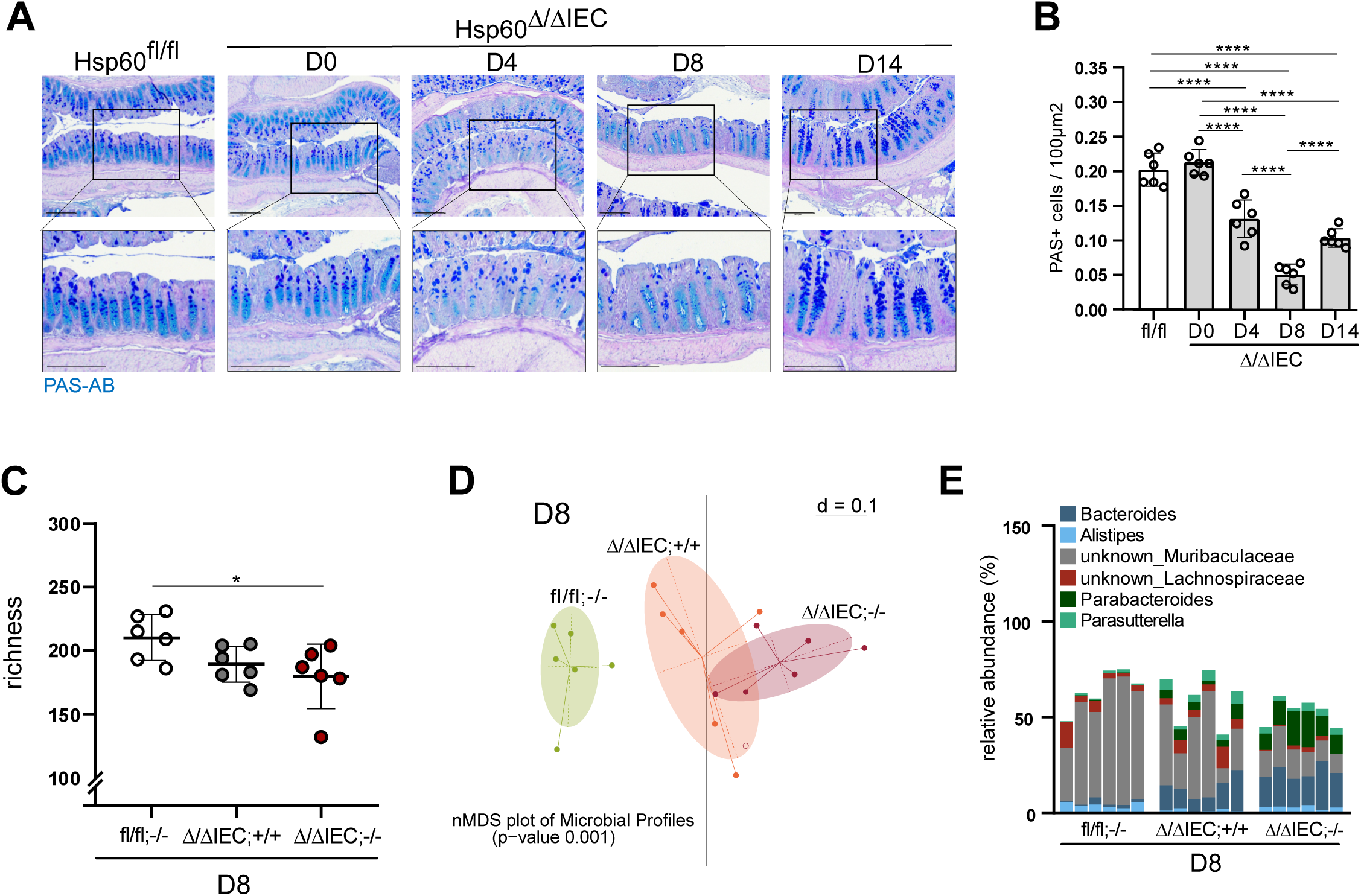
Metabolic injury drives dysbiotic changes in the intestinal microbiota. Related to Figure 2. **A,** Representative images of PAS-AB staining in colonic tissue of Hsp60^Δ/ΔIEC^ and Hsp60^fl/fl^ mice from day 0 to 14. Acidic mucins are stained in cyan, neutral mucins in dark blue. Nuclei are stained with hematoxylin. Scale bars: 200µm. **B,** Quantification of PAS positive cells per 100µm^2^ in colonic tissue of Hsp60^Δ/ΔIEC^ and Hsp60^fl/fl^ from day 0 to 14. **C,** Species richness in colonic content of Hsp60^Δ/ΔIEC^;Il10^-/-^ (Ctrl: Hsp60^Δ/ΔIEC^;Il10^+/+^, Hsp60^fl/fl^;Il10^-/-^) mice at day 8. **D,** β-diversity depicted as nMDS plot of microbial profiles of colonic content stratified by genotype (Hsp60^Δ/ΔIEC^;Il10^-/-^, Hsp60^Δ/ΔIEC^;Il10^+/+^ and Hsp60^fl/fl^;Il10^-/-^ mice) at day 8. **E,** Relative abundance of significantly enriched genera in Hsp60^Δ/ΔIEC^;Il10^-/-^ (Ctrl: Hsp60^Δ/ΔIEC^;Il10^+/+^, Hsp60^fl/fl^;Il10^-/-^) mice at day 8. (**B, C**) Data are represented by mean ± s.d. of six biological replicates. P-values were calculated by ANOVA followed by pairwise comparison testing (Tukey). *P<0.05, **P<0.01, ***P<0.001, ****P<0.0001. (**E**) Kruskal-Wallis test was used to select significant genera (P<0.05). D: day; Hsp60: heat shock protein 60; IECs: intestinal epithelial cells; IL10: interleukin-10; PAS-AB: Alcian blue/Periodic acid-Schiff.

**Figure S4:**
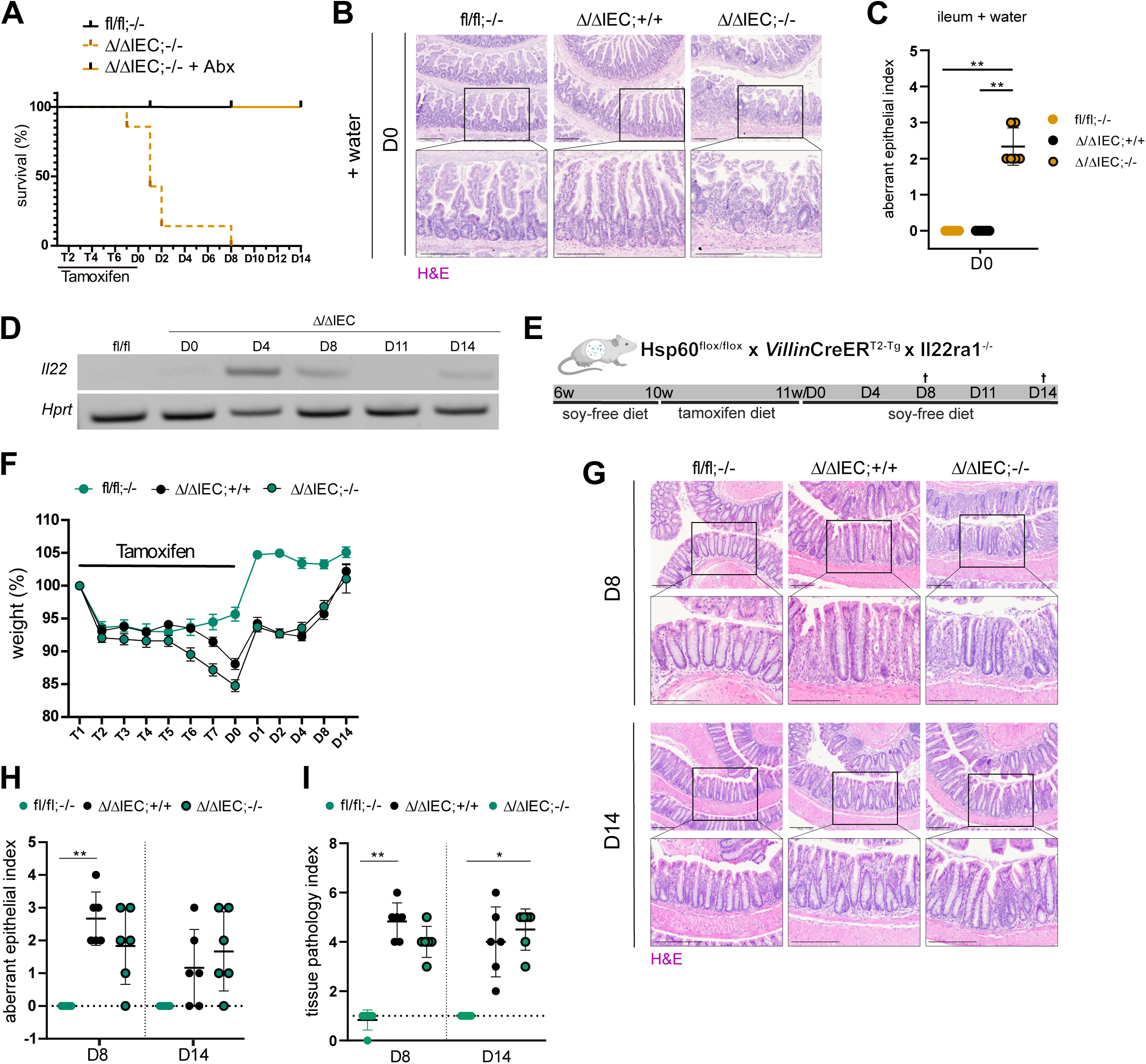
AhR protective mechanisms on intestinal homeostasis are independent of IL-22-signaling. Related to Figure 3. **A,** Survival (%) of Hsp60^Δ/ΔIEC^;AhR^-/-^ mice treated with or without Abx (Ctrl: Hsp60^fl/fl^;AhR^-/-^) from start of tamoxifen treatment until day 14. **B, C,** (**B**) Representative images of H&E staining and (**C**) histological scoring for an aberrant epithelium of ileal tissue of Hsp60^Δ/ΔIEC^;AhR^-/-^ (Ctrl: Hsp60^Δ/ΔIEC^;AhR^+/+^, Hsp60^fl/fl^;AhR^-/-^) mice at day 0 treated without Abx. Scale bars: 200µm. **D,** Agarose gel of qPCR from *Il22* (and the housekeeper *Hprt*) using mRNA of whole colonic tissue from day 0 to 14 in Hsp60^Δ/ΔIEC^ and Hsp60^fl/fl^ mice. One gel line includes a mix of six biological replicates. **E,** Experimental setup. Hsp60^fl/fl^ x *Villin*CreER^T2-Tg^ x Il22ra1^-/-^ mice (Hsp60^Δ/ΔIEC^;Il22ra1^-/-^; Ctrl: Hsp60^Δ/ΔIEC^;Il22ra1^+/+^, Hsp60^fl/fl^;Il22ra1^-/-^) were fed a soy-free diet for 4 weeks, followed by a tamoxifen diet at the age of 10 weeks for 7 days to induce Hsp60 deletion. Mice were sampled at day 8 and 14 day after tamoxifen feeding and full Hsp60 deletion. **F,** Relative weight curve of Hsp60^Δ/ΔIEC^;Il22ra1^-/-^ mice (Ctrl: Hsp60^Δ/ΔIEC^;Il22ra1^+/+^, Hsp60^fl/fl^;Il22ra1^-/-^) from the start of tamoxifen feeding until sampling time point. **G, H, I,** (**G**) Representative images of H&E staining and (**H**) histological scoring for an aberrant epithelium and (**I**) inflammation of distal colonic tissue of Hsp60^Δ/ΔIEC^;Il22ra1^-/-^ mice (Ctrl: Hsp60^Δ/ΔIEC^;Il22ra1^+/+^, Hsp60^fl/fl^;Il22ra1^-/-^) at day 8 and 14. Scale bars: 200µm. Data are represented by (**F**) mean ± sem or (**A, C, H, I**) mean ± s.d. of six biological replicates. (**C, H, I**) P-values were calculated by Kruskal-Wallis test followed by pairwise comparison testing (Dunn). *P<0.05, **P<0.01, ***P<0.001, ****P<0.0001. Hsp60: heat shock protein 60; IECs: intestinal epithelial cells; AhR: aryl hydrocarbon receptor; Abx: Antibiotics; H&E: hematoxylin and eosin; IL-22: interleukin 22; Il22ra1: interleukin 22 receptor subunit-α1.

**Figure S5:**
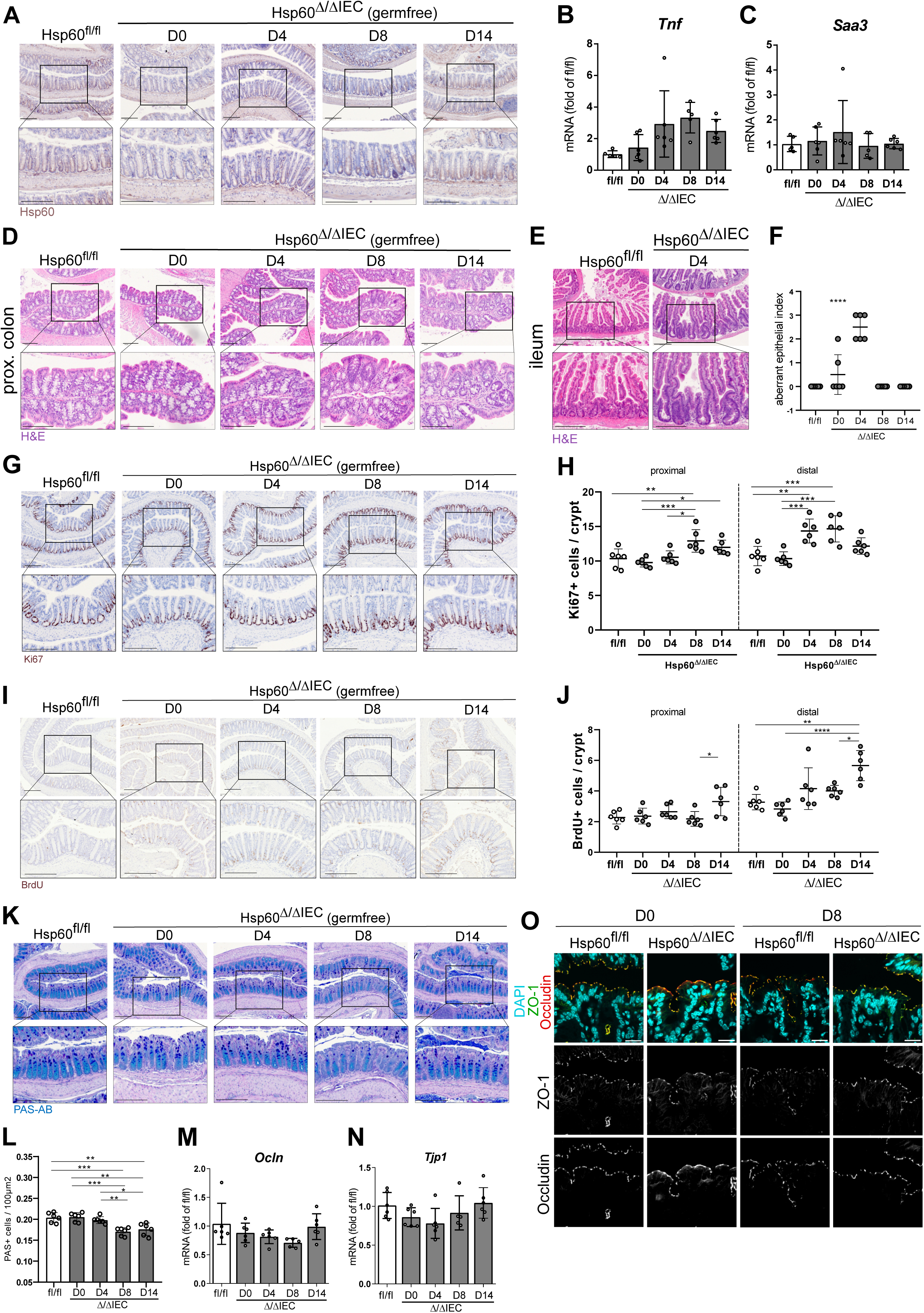
The epithelial barrier is not affected by mitochondrial dysfunction under germ-free conditions. Related to Figure 4. **A,** Representative images of immunohistochemistry staining of Hsp60 in distal colonic tissue of GF Hsp60^Δ/ΔIEC^ and Hsp60^fl/fl^ mice from day 0 to14. Brown color shows specific Hsp60 signal, nuclei are stained with hematoxylin. Scale bars: 200µm. **B, C,** qPCR of (**B**) *Tnf* and (**C**) *Saa3* using mRNA of whole colonic tissue from day 0 to 14 of GF Hsp60^Δ/ΔIEC^ (Ctrl: Hsp60^fl/fl^) mice. **D, E,** Representative images of H&E staining from (**D**) day 0 to 14 of proximal colonic and (**E**) day 4 of ileal tissue of GF Hsp60^Δ/ΔIEC^ and Hsp60^fl/fl^ mice. Scale bars: 200µm. **F,** H&E stainings of ileal tissue of GF Hsp60^Δ/ΔIEC^ mice from day 0 to 14 were histologically scored for an aberrant epithelium. **G, I**, Representative images of immunohistochemistry staining of the proliferation markers (**G**) Ki67 and (**I**) incorporated BrdU two hours after intraperitoneal injection in distal colonic tissue of Hsp60^Δ/ΔIEC^ and Hsp60^fl/fl^ mice from day 0 to 14. Brown color shows specific Ki67 or BrdU signal, nuclei are stained with hematoxylin. Scale bars: 200µm. **H, J**, Quantification of (**H**) Ki67-positive cells per crypt and (**J**) incorporated BrdU-positive cells per crypt in colonic tissue of Hsp60^Δ/ΔIEC^ and Hsp60^fl/fl^ mice from day 0 to 14. **K**, Representative images of PAS-AB staining in distal colonic tissue of GF Hsp60^Δ/ΔIEC^ and Hsp60^fl/fl^ mice from day 0 to 14. Acidic mucins are stained in cyan, neutral mucins in dark blue. Nuclei are stained with hematoxylin. Scale bars: 200µm. **L,** Quantification of PAS positive cells per 100µm^2^ in distal colonic tissue of GF Hsp60^Δ/ΔIEC^ and Hsp60^fl/fl^ from day 0 to 14. **M, N,** qPCR of tight junctional protein genes (**M**) *Ocln* and (**N**) *Tjp1* using mRNA of whole colonic tissue from day 0 to 14 in GF Hsp60^Δ/ΔIEC^ and Hsp60^fl/fl^ mice. **O**, Representative immunofluorescent co-staining of tight junction proteins ZO-1 (green) and Occludin (red) as well as DAPI (blue) at day 0 and 8 in the surface epithelium in distal colonic tissue of GF Hsp60^Δ/ΔIEC^ and Hsp60^fl/fl^ mice. Scale bars: 20µm. Data are represented by mean ± s.d. of six biological replicates. (**B, C, F, L**) P-values were calculated by One-way ANOVA followed by pairwise comparison testing (Tukey). (**H, J, M, N**) P-values were calculated by Kruskal-Wallis test followed by pairwise comparison testing (Dunn). *P<0.05, **P<0.01, ***P<0.001, ****P<0.0001. D: day; GF: germfree; Hsp60: heat shock protein 60; IECs: intestinal epithelial cells; H&E: hematoxylin and eosin; BrdU: Bromodeoxyuridine; Ocln: Occludin; Tjp1: tight junction protein 1; ZO-1: Zonula Occludens-1; PAS-AB: Alcian blue/Periodicacid-Schiff; DAPI: 4′,6-diamidino-2-phenylindole.

**Figure S6:**
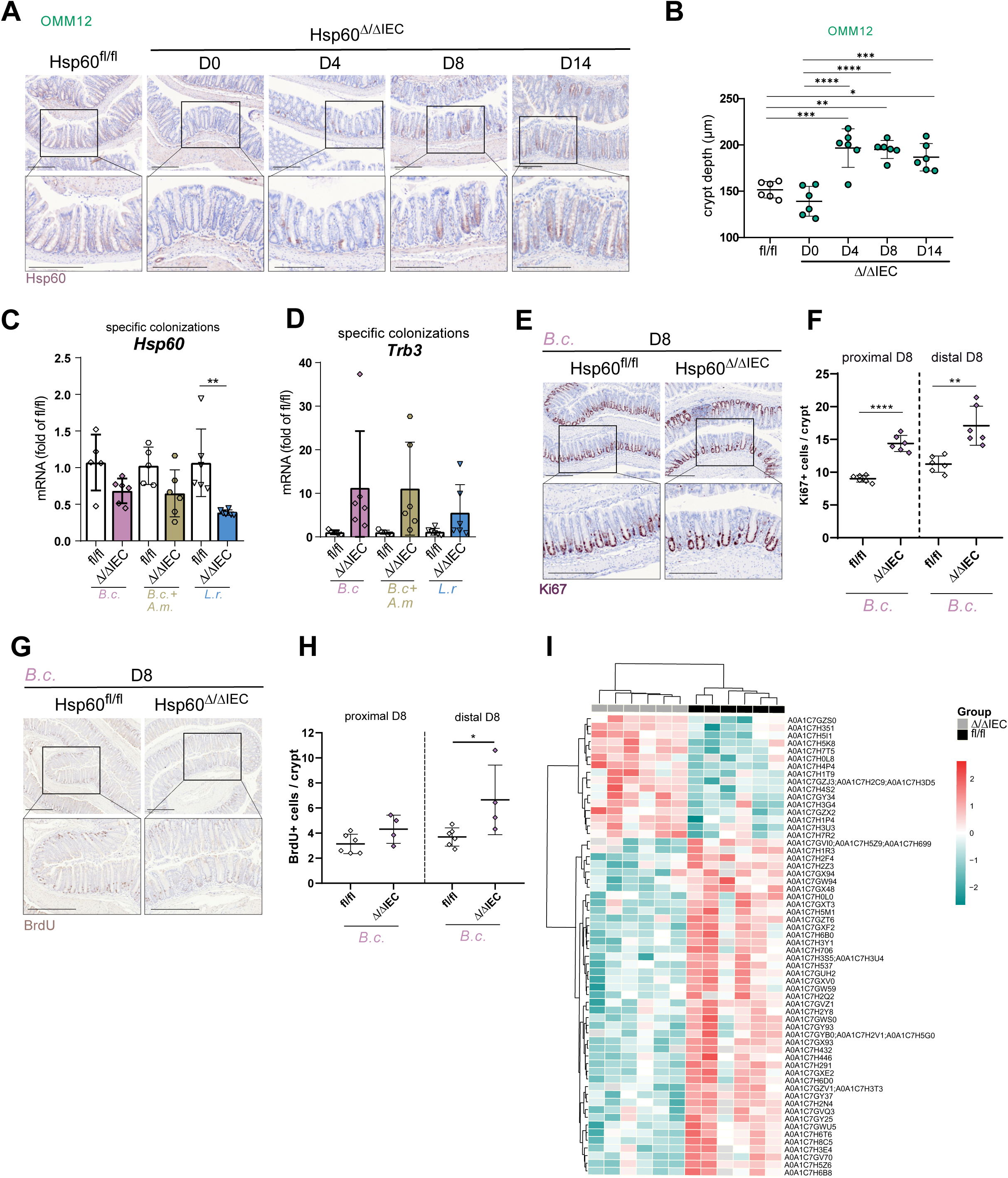
*Bacteroides caecimuris* induces metabolic injury. Related to Figure 5. **A,** Representative images of immunohistochemistry staining of Hsp60 of distal colonic tissue of GF Hsp60^Δ/ΔIEC^ and Hsp60^fl/fl^ mice colonized with the Oligo-Mouse-Microbiota (OMM^12^) synthetic bacterial community from day 0 to 14 after Hsp60 deletion. Scale bars: 200µm. **B,** Crypt depth of H&E stainings of distal colonic tissue of GF Hsp60^Δ/ΔIEC^ and Hsp60^fl/fl^ mice colonized with OMM^12^ was measured from day 0 to 14. **C, D,** qPCR of (**C**) the mitochondrial chaperone *Hsp60* and (**D**) the MT-UPR marker gene *Trb3* using mRNA of whole colonic tissue of GF Hsp60^Δ/ΔIEC^ and Hsp60^fl/fl^ mice colonized with *Bacteroides caecimuris*, *Bacteroides caecimuris* and *Akkermanisa muciniphila* or *Limosilactobacillus reuteri* sampled on day 8. **E,** Representative images of immunohistochemistry staining of the proliferation marker Ki67 in distal colonic tissue of GF Hsp60^Δ/ΔIEC^ and Hsp60^fl/fl^ mice colonized with *B. caecimuris* on day 8. Brown color shows specific Ki67 signal, nuclei are stained with hematoxylin. Scale bars: 200µm. **F,** Quantification of Ki67-positive cells per crypt in GF Hsp60^Δ/ΔIEC^ and Hsp60^fl/fl^ mice colonized with *B. caecimuris* on day 8. **G,** Representative images of immunohistochemistry staining of incorporated BrdU in distal colonic tissue of GF Hsp60^Δ/ΔIEC^ and Hsp60^fl/fl^ mice colonized with *B. caecimuris* on day 8 two hours after intraperitoneal injection. Brown color shows specific BrdU signal, nuclei are stained with hematoxylin. Scale bars: 200µm. **H,** Quantification of BrdU-positive cells per crypt in GF Hsp60^Δ/ΔIEC^ and Hsp60^fl/fl^ mice colonized with *B. caecimuris* on day 8. **I,** Heatmap of normalized (z-score) protein intensities (of identified protein groups assigned to *B. caecimuris* and differentially abundant in mono-colonized-GF Hsp60^Δ/ΔIEC^ and Hsp60^fl/fl^ mice with *B. caecimuris*, based on non-corrected P<0.01). Non-detected protein groups (protein intensities with “NA” in Supplementary material table 1) are colored in gray. Data are represented by mean ± s.d. of six biological replicates (**B**, **C, D, F, H**). P-values were calculated by ANOVA followed by pairwise comparison testing (Tukey). *P<0.05, **P<0.01, ***P<0.001, ****P<0.0001. GF: germfree; OMM^12^: Oligo-Mouse-Microbiota 12; D: day; Hsp60: heat shock protein 60; IECs: intestinal epithelial cells; MT-UPR: mitochondrial unfolded protein response; H&E: hematoxylin and eosin; B.c: *Bacteroides caecimuris*; A.m: *Akkermansia muciniphila*; L.r: *Limosilactobacillus reuteri*; Trb3: tribbles pseudokinase 3; BrdU: Bromodeoxyuridine.

**Figure S7:**
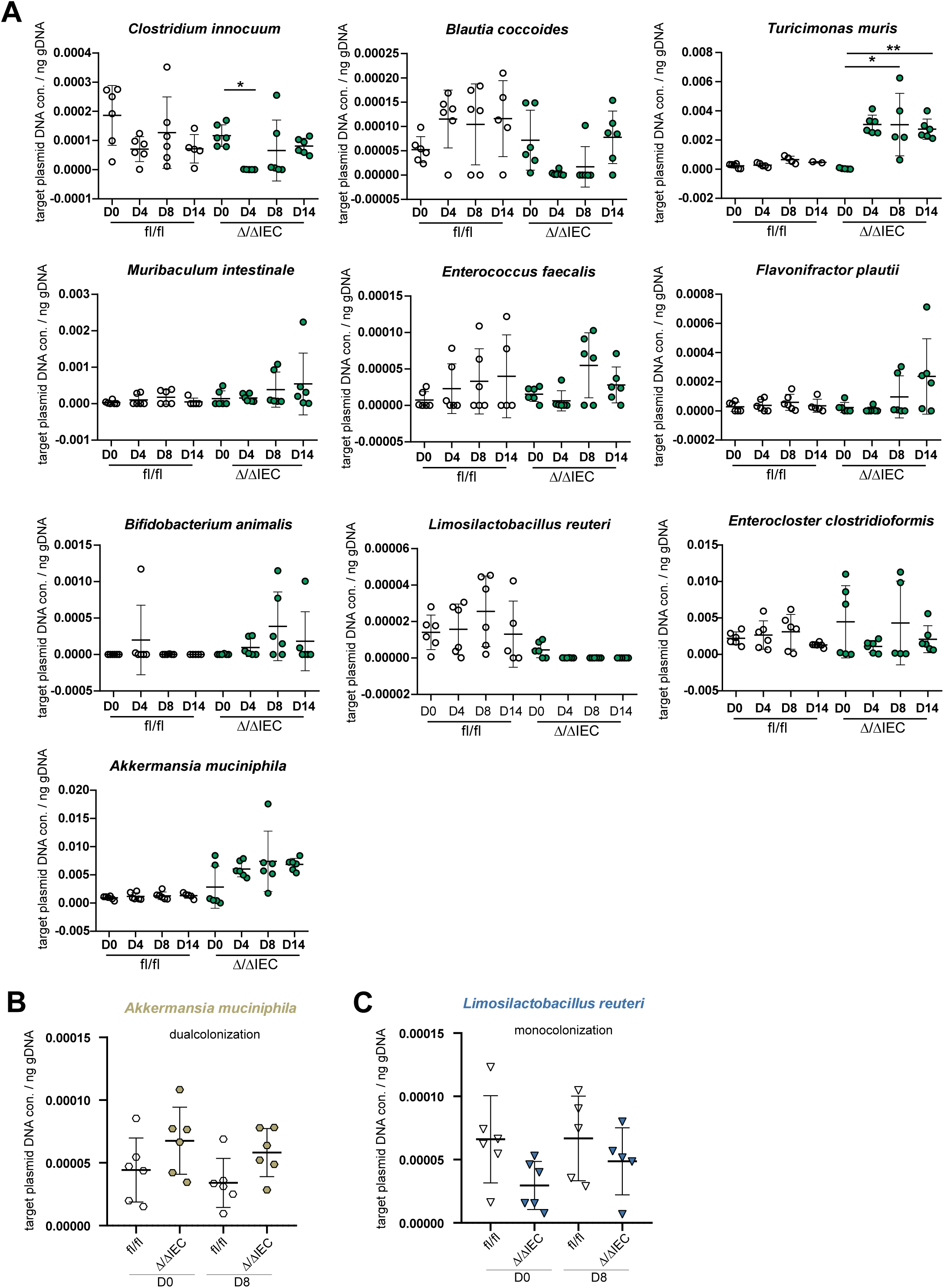
Relative abundance of bacteria in selective colonizations. Related to Figure 5. **A,** Abundance of 10 different species from the Oligo-Mouse-Microbiota (OMM^12^) synthetic bacterial community in colonic content of GF Hsp60^Δ/ΔIEC^ and Hsp60^fl/fl^ mice colonized with OMM^12^ detected by 16S rRNA targeted qPCR from day 0 to 14 (*Acutalibacter muris* signal was below detection limits). **B,** Abundance of *Akkermanisa muciniphila* in colonic content of GF Hsp60^Δ/ΔIEC^ and Hsp60^fl/fl^ mice dual-colonized with *Bacteroides caecimuris* and *Akkermanisa muciniphila* detected by 16S rRNA targeted qPCR at day 0 and 8. **C,** Relative abundance of *Limosilactobacillus reuteri* in colonic content of GF Hsp60^Δ/ΔIEC^ and Hsp60^fl/fl^ mice colonized with *Limosilactobacillus reuteri* detected by 16S rRNA targeted qPCR at day 0 and 8. Five to six biological replicates were measured and data are represented by mean ± s.d. of two to six biological replicates depending on detection. (**A**) P-values were calculated by Kruskal-Wallis test followed by pairwise comparison testing (Dunn). (**B, C**) P-values were calculated by ANOVA followed by pairwise comparison testing (Tukey). *P<0.05, **P<0.01, ***P<0.001, ****P<0.0001. GF: germfree; D: day; Hsp60: heat shock protein 60; IECs: intestinal epithelial cells.

**Figure S8:**
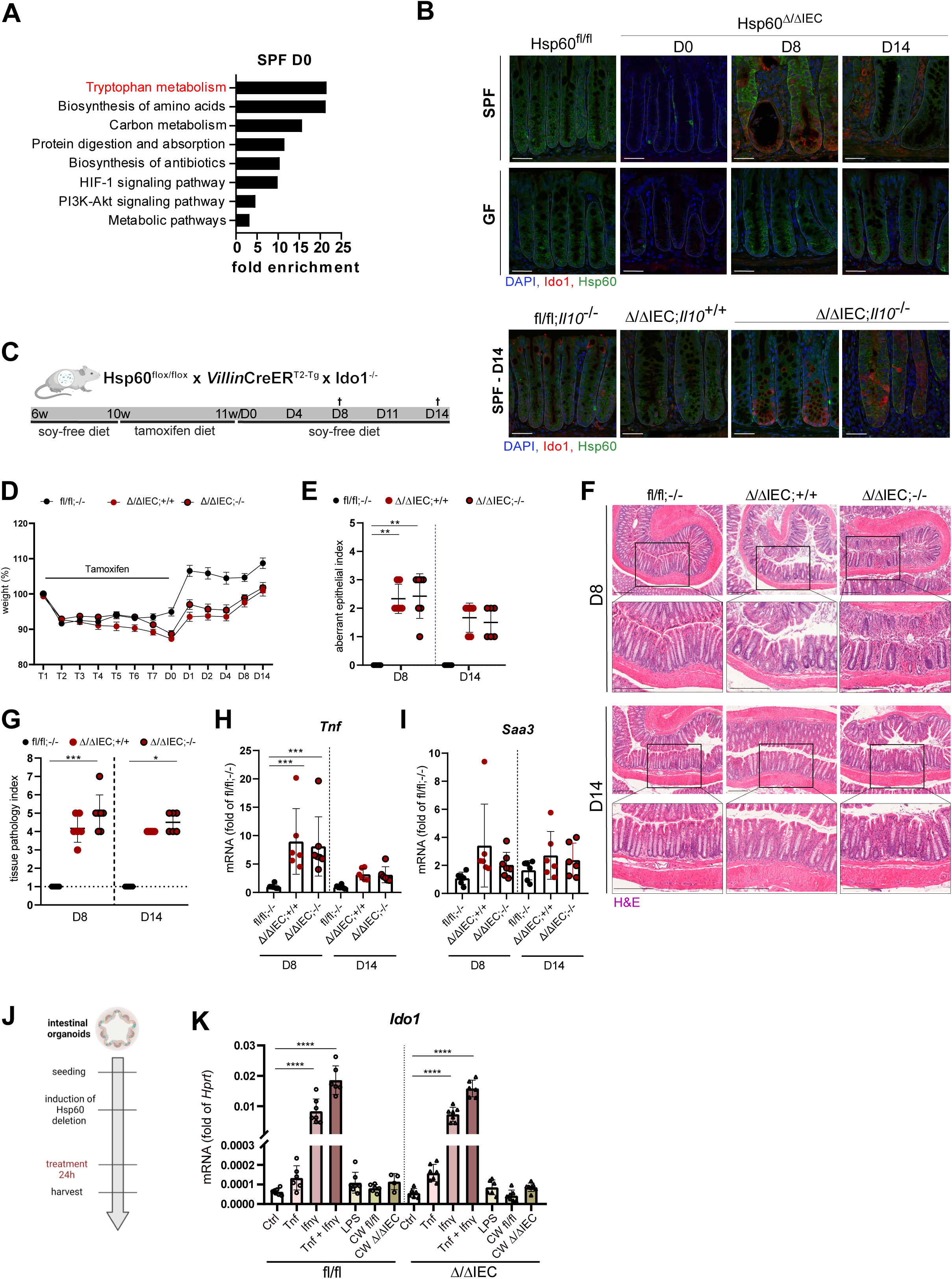
Host-tryptophan metabolism is rewired during metabolic injury. Related to Figure 6. **A,** KEGG pathway analysis of the 45 altered genes in Hsp60^Δ/ΔIEC^ mice under SPF conditions at day 0. Only pathways with a significant False Discovery Rate and at least 5 genes mapping to a pathway were selected. **B,** Representative immunofluorescent co-staining of Hsp60 (green), Ido1 (red), and DAPI (blue) in distal colonic tissue. Upper panel: Hsp60^Δ/ΔIEC^ and Hsp60^fl/fl^ mice under SPF or GF housing from day 0 to 14. Lower panel: Hsp60^Δ/ΔIEC^;Il10^-/-^ (Ctrl: Hsp60^Δ/ΔIEC^;Il10^+/+^, Hsp60^fl/fl^;Il10^-/-^) mice under SPF housing at day 14. Scale bars: 50 µm. **C,** Experimental setup. Hsp60^fl/fl^ x *Villin*CreER^T2-Tg^ x Ido1^-/-^ mice (Hsp60^Δ/ΔIEC^;Ido1^-/-^; Ctrl: Hsp60^Δ/ΔIEC^;Ido1^+/+^, Hsp60^fl/fl^;Ido1^-/-^) were fed a soy-free diet for 4 weeks, followed by a tamoxifen diet at the age of 10 weeks for 7 days to induce Hsp60 deletion. Mice were sampled at day 8 and 14 after tamoxifen feeding and full Hsp60 deletion. **D,** Relative weight curve of Hsp60^Δ/ΔIEC^;Ido1^-/-^ mice (Ctrl: Hsp60^Δ/ΔIEC^;Ido1^+/+^, Hsp60^fl/fl^;Ido1^-/-^) from the start of tamoxifen feeding until sampling time point. **E, F, G,** (**E**) Representative images of H&E staining and (**F**) histological scoring for an aberrant epithelium and (**G**) inflammation of distal colonic tissue of Hsp60^Δ/ΔIEC^;Ido1^-/-^ mice (Ctrl: Hsp60^Δ/ΔIEC^;Ido1^+/+^, Hsp60^fl/fl^;Ido1^-/-^) at day 8 and 14. Scale bars: 200µm. **H, I,** qPCR of the inflammatory marker genes (**H**) *Tnf* and (**I**) *Saa3* using mRNA of whole colonic tissue on day 8 and 14 in Hsp60^Δ/ΔIEC^;Ido1^-/-^ mice (Ctrl: Hsp60^Δ/ΔIEC^;Ido1^+/+^, Hsp60^fl/fl^;Ido1^-/-^). **J,** Experimental setup. Small intestinal crypts were isolated and seeded in Matrigel^®^, followed by Hsp60 deletion *ex vivo* via tamoxifen addition to the media. Organoids were treated 5 days after Hsp60 deletion for 24h. **K,** qPCR of *Ido1* using mRNA of organoids at day 6 after Hsp60 deletion that were treated with Tnf, Ifnγ, LPS or cecal water (from Hsp60^Δ/ΔIEC^ or Hsp60^fl/fl^ mice at day 8) for 24h. Data are represented by (**D**) mean ± sem or (**E, G-I, K**) mean ± s.d. of six biological replicates. P-values were calculated by (**E, G**) Kruskal-Wallis test followed by pairwise comparison testing (Dunn) or by (**H, I, K**) ANOVA followed by pairwise comparison testing (Tukey). *P<0.05, **P<0.01, ***P<0.001, ****P<0.0001. D: day; Hsp60: heat shock protein 60; IECs: intestinal epithelial cells; Ido1: indoleamine 2, 3-dioxygenase 1; SPF: specific pathogen-free; GF: germ-free; DAPI: 4′,6-diamidino-2-phenylindole; Tnf: tumor necrosis factor; Saa3: serum amyloid A 3; H&E: hematoxylin and eosin; Ifnγ: interferon γ; LPS: lipopolysaccharide; CW: cecal water.

**Figure S9:**
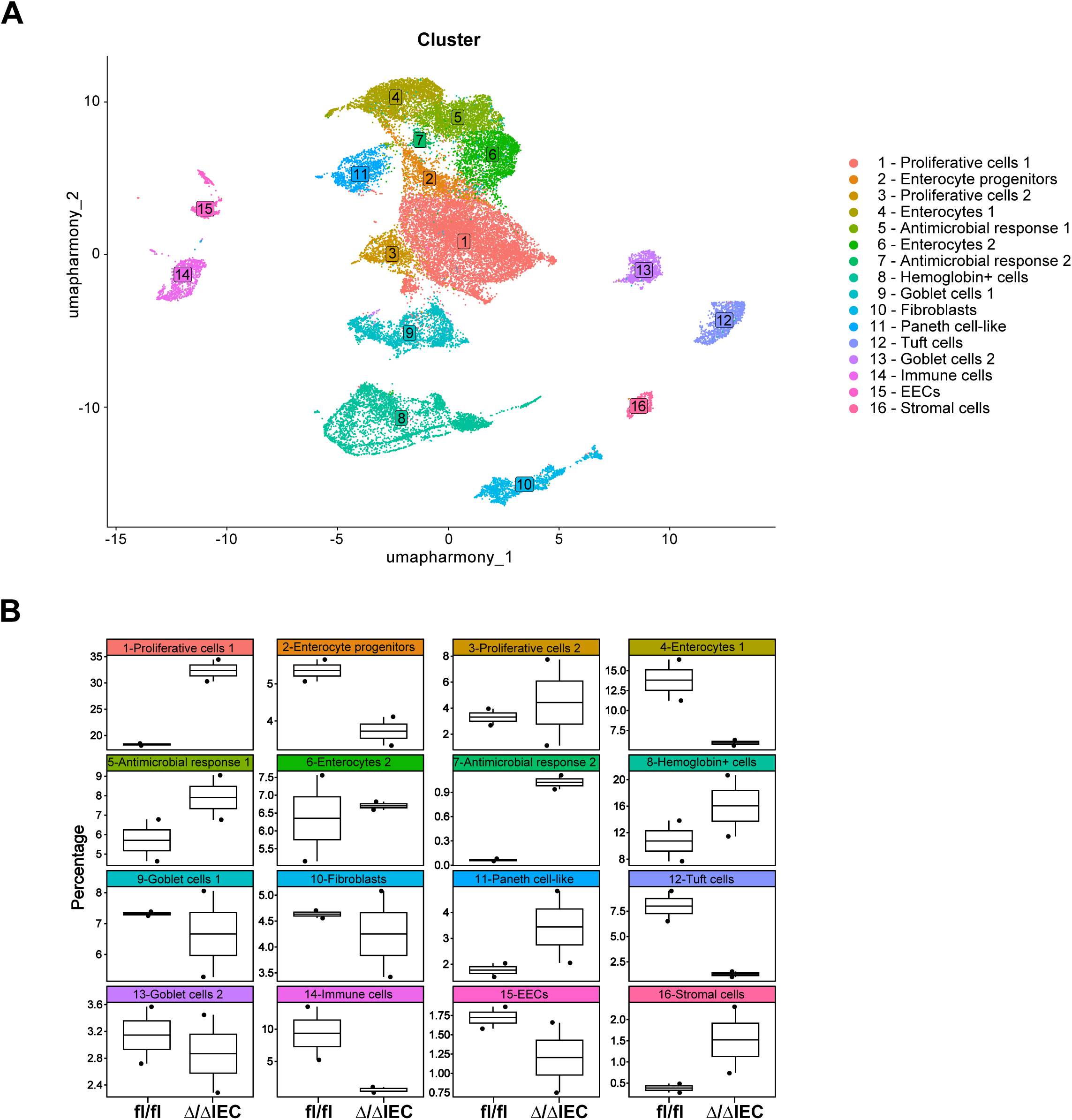
Single-cell resolution of metabolically-injured colonic cells. Related to Figure 6. **A,** UMAP visualization of all cell clusters colored by cluster identity across all conditions. Colonic epithelial cells were isolated from Hsp60^Δ/ΔIEC^ (n=2) and Hsp60^fl/fl^ mice (n=2) at day 8 post-tamoxifen. Note the presence of residual non-epithelial cells. DEGs defining clusters are available in Supplementary table 2. **B,** Distribution (in %) of cells in each cluster and per condition (n=2 for Hsp60^Δ/ΔIEC^ mice and n=2 for Hsp60^fl/fl^ mice). Data include two biological replicates per condition (**A, B**). UMAP: Uniform manifold approximation and projection; IECs: intestinal epithelial cells; EECs: enteroendocrine cells; DEGs: differentially expressed genes.

**Supplementary Table 1:**
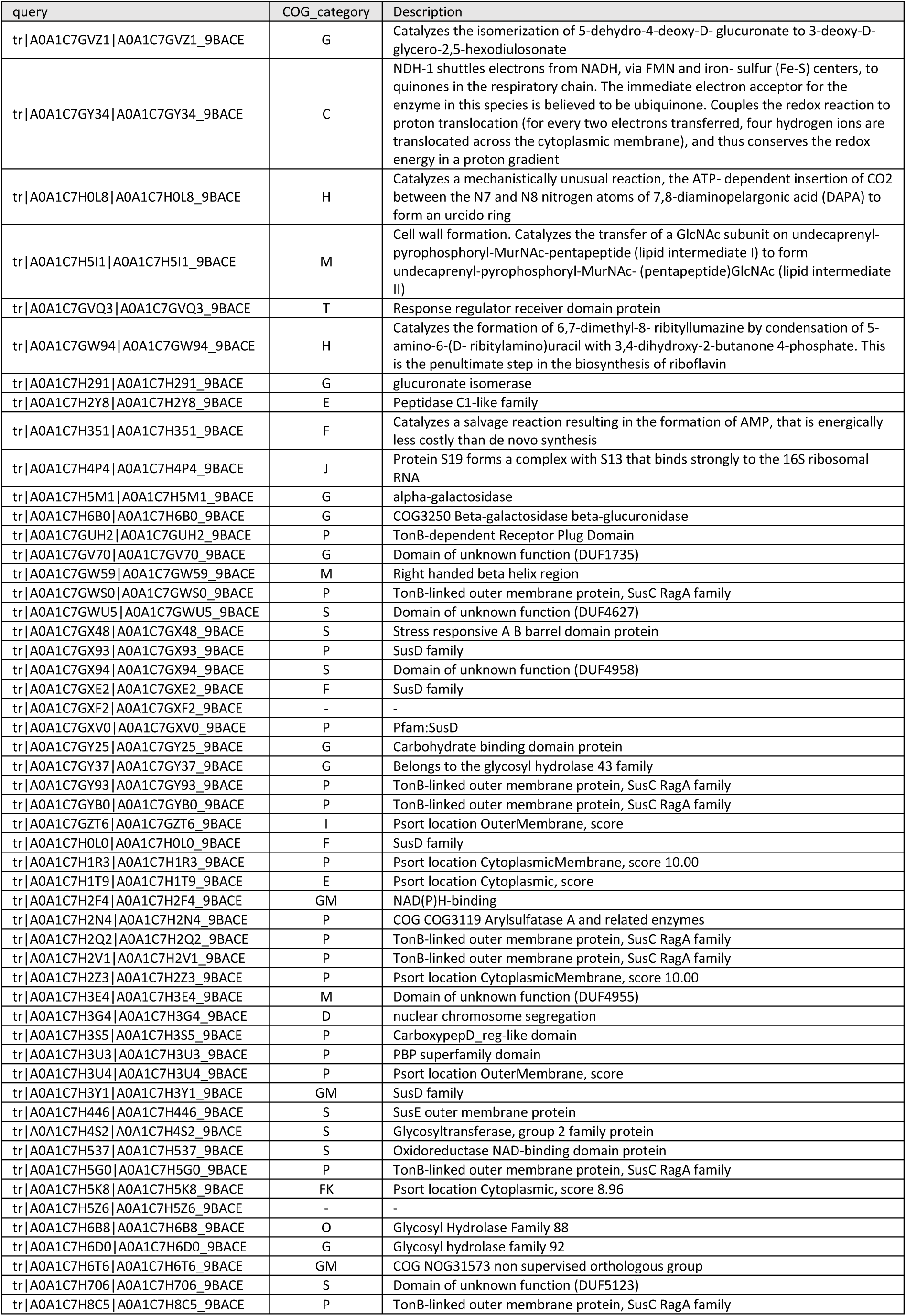
Mapping of Protein-Groups to Clusters of Orthologous Genes (COGs). Related to Figure 5.

**Supplementary Table 3:** Mouse models included in the study. Related to STAR Methods.

| IEC-specific inducible knock-out (KO) |  | Nomenclature |
| --- | --- | --- |
| Hsp60<sup>flox/flox</sup> x *Villin*CreER<sup>T2-Tg</sup><br>(control: Hsp60<sup>flox/flox</sup> x *Villin*CreER<sup>T2-wt</sup>) |  | Hsp60<sup>Δ/ΔIEC</sup><br>(control: Hsp60<sup>fl/fl</sup>) |
| IEC-specific inducible KO | Constitutive whole body KO | Nomenclature |
| Hsp60<sup>flox/flox</sup> x *Villin*CreER<sup>T2-Tg</sup><br>(control: Hsp60<sup>flox/flox</sup> x *Villin*CreER<sup>T2-wt</sup>) | Interleukin-10 (*Il10*<sup>-/-</sup>)<br>(control: *Il10*<sup>+/+</sup>) | Hsp60<sup>Δ/ΔIEC</sup>; *Il10*<sup>-/-</sup><br>(control: Hsp60<sup>fl/fl</sup>; *Il10*<sup>-/-</sup>,<br>Hsp60<sup>Δ/ΔIEC</sup>; *Il10*<sup>+/+</sup>) |
|  | Aryl hydrocarbon receptor (*AhR*<sup>-/-</sup>)<br>(control: *AhR*<sup>+/+</sup>) | Hsp60<sup>Δ/ΔIEC</sup>; *AhR*<sup>-/-</sup><br>(control: Hsp60<sup>fl/fl</sup>; *AhR*<sup>+/+</sup>,<br>Hsp60<sup>fl/fl</sup>; *AhR*<sup>-/-</sup>,<br>Hsp60<sup>Δ/ΔIEC</sup>; *AhR*<sup>+/+</sup>) |
|  | Interleukin-22 receptor subunit-α1 (*Il22ra1*<sup>-/-</sup>) (control: *Il22ra1*<sup>+/+</sup>) | Hsp60<sup>Δ/ΔIEC</sup>; *Il22ra1*<sup>-/-</sup><br>(control: Hsp60<sup>fl/fl</sup>; *Il22ra1*<sup>-/-</sup>,<br>Hsp60<sup>Δ/ΔIEC</sup>; *Il22ra1*<sup>+/+</sup>) |
|  | Indoleamine 2,3-dioxygenase 1 (*Ido1*<sup>-/-</sup>)<br>(control: *Ido1*<sup>+/+</sup>) | Hsp60<sup>Δ/ΔIEC</sup>; *Ido1*<sup>-/-</sup><br>(control: Hsp60<sup>fl/fl</sup>; *Ido1*<sup>-/-</sup>,<br>Hsp60<sup>Δ/ΔIEC</sup>; *Ido1*<sup>+/+</sup>) |
| IEC-specific inducible overexpression |  | Nomenclature |
| Atf5<sup>flox/flox</sup> x *Villin*CreER<sup>T2-Tg</sup><br>(control: Atf5<sup>flox/flox</sup> x *Villin*CreER<sup>T2-wt</sup>) |  | Atf5<sup>IEC</sup><br>(control: Atf5<sup>fl/fl</sup>) |

**Supplementary Table 2: Differentially expressed genes in all Louvain clusters and functional annotation of cluster 1, 5, 7 and 16, respectively. Related to Figure 6 and Figure S9.**

